# Extensive genome evolution distinguishes maize within a stable tribe of grasses

**DOI:** 10.1101/2025.01.22.633974

**Authors:** Michelle C. Stitzer, Arun S. Seetharam, Armin Scheben, Sheng-Kai Hsu, Aimee J. Schulz, Taylor M. AuBuchon-Elder, Mohamed El-Walid, Taylor H. Ferebee, Charles O. Hale, Thuy La, Zong-Yan Liu, Sarah J. McMorrow, Patrick Minx, Alyssa R. Phillips, Michael L. Syring, Travis Wrightsman, Jingjing Zhai, Rémy Pasquet, Christine A. McAllister, Simon T. Malcomber, Paweena Traiperm, Daniel J. Layton, Jinshun Zhong, Denise E. Costich, R. Kelly Dawe, Kevin Fengler, Charlotte Harris, Zach Irelan, Victor Llaca, Praveena Parakkal, Gina Zastrow-Hayes, Margaret R. Woodhouse, Ethalinda K. Cannon, John L. Portwood, Carson M. Andorf, Patrice S. Albert, James A. Birchler, Adam Siepel, Jeffrey Ross-Ibarra, M. Cinta Romay, Elizabeth A. Kellogg, Edward S. Buckler, Matthew B. Hufford

## Abstract

Over the last 20 million years, the Andropogoneae tribe of grasses has evolved to dominate 17% of global land area. Domestication of these grasses in the last 10,000 years has yielded our most productive crops, including maize, sugarcane, and sorghum. The majority of Andropogoneae species, including maize, show a history of polyploidy – a condition that, while offering the evolutionary advantage of multiple gene copies, poses challenges to basic cellular processes, gene expression, and epigenetic regulation. Genomic studies of polyploidy have been limited by sparse sampling of taxa in groups with multiple polyploidy events. Here, we present 33 genome assemblies from 27 species, including chromosome-scale assemblies of maize relatives *Zea* and *Tripsacum*. In maize, the after-effects of polyploidy have been widely studied, showing reduced chromosome number, biased fractionation of duplicate genes, and transposable element (TE) expansions. While we observe these patterns within the genus *Zea*, 12 other polyploidy events deviate significantly. Those tetraploids and hexaploids retain elevated chromosome number, maintain nearly complete complements of duplicate genes, and have only stochastic TE amplifications. These genomes reveal variable outcomes of polyploidy, challenging simple predictions and providing a foundation for understanding its evolutionary implications in an ecologically and economically important clade.

## Introduction

Andropogoneae grasses have been integral to the origins and spread of grasslands, relevant to human culture and agriculture for thousands of years, and are of growing importance for their ability to rebalance the carbon cycle. These 1,200 species of warm-season grasses dominate ecosystems in Africa, North and South America, and portions of South and Southeast Asia (Gibson, 2009; Kellogg, 2015). The ancestor of these grasses evolved the highly efficient C4 photosynthesis (Bianconi et al., 2020), and all are adapted for low or variable CO_2_ abundance, meaning these species are nitrogen and water use efficient (Ghannoum et al., 2011; Morison & Gifford, 1983; Rawson et al., 1977). Today, these grasses include the largest production crops maize and sugarcane, drought resistant sorghum, the bioenergy crop *Miscanthus*, and numerous forage grasses. Altogether, Andropogoneae grasses cover 17% of global land (Lehmann et al., 2019).

Allopolyploidy has been a major force in the evolution of Andropogoneae grasses, with at least ⅓ of all speciation events associated with polyploidy (Estep et al., 2014). Newly formed polyploids often face meiotic abnormalities and establishment challenges (Ramsey & Schemske, 2002), and genome doubling can induce epigenetic instability (Comai, 2005). Considering these challenges, it is surprising there are so many polyploids, not only in the Andropogoneae, but across the plant kingdom (Alix et al., 2017). Explanations for their prevalence range from ecological novelty of polyploids, allowing them to exploit niches or stressful environments unavailable to their diploid progenitors (Van de Peer et al., 2017), to the increased fitness and adaptive potential arising from the permanent hybridity of polyploids (Roose & Gottlieb, 1976; Stebbins, 1959). Over longer time scales, polyploids frequently revert to a diploid-like state (Baduel et al., 2018; Comai, 2005; Leitch & Leitch, 2008; Otto, 2007; J. F. Wendel, 2000, 2015), via three oft-cited mechanisms - chromosomal reduction, gene loss via fractionation, and transposable element amplification.

Despite these general patterns, there are major gaps in our understanding of even the best known polyploid systems (Soltis et al., 2016). Mounting evidence suggests that at least some polyploid lineages undergo minimal genome evolution (cotton, Wendel & Cronn, 2003, bamboo, Ma et al., 2024; *Arabidopsis suecica,* Burns et al., 2021), and other plant lineages rarely produce polyploids (gymnosperms, Ickert-Bond et al., 2020), illustrating exceptions to the conventional models. Here, we present genome assemblies of 33 Andropogoneae individuals from 27 species, capturing 14 independent polyploid formation events. Of these, twelve have little evidence of large-scale genomic reorganization, show unpredictable TE dynamics, and retain multiples of the base chromosome number and most genes. In contrast, the lineage leading to *Zea* exhibits notable changes, with expanded TEs, and only retaining a subset of progenitor genes in a radically rearranged karyotype.

## Results and Discussion

### Genome assemblies of Andropogoneae

We generated highly contiguous assemblies of 33 members of Andropogoneae, representing 27 species (Supplemental Table S1). Sequencing technologies evolved throughout our project, so while most individuals were sequenced with PacBio HiFi (27 plants), 5 plants were sequenced with PacBio Continuous Long Read (CLR) sequencing, and one with Oxford Nanopore Technology (ONT) (Table S2). To further improve contiguity, 24 assemblies were scaffolded using BioNano optical maps, HiC, or genetic maps (Table S2). Most samples come from outbred accessions, and many come from species with multiple ploidy levels and mixed-ploidy populations.

Our assemblies include nine chromosome-scale assemblies of all diploid (2n=20) teosinte species and subspecies within *Zea*, as well as two individuals of *Tripsacum dactyloides* (2n=36), together representing the subtribe Tripsacinae. The 22 additional species sequenced within the tribe Andropogoneae (Supplemental Table S1) represent major clades in the phylogeny (Figure 1A) and the global distribution (Figure 1C). We observed a broad range of haploid assembly sizes, from 663 Mb to 4,767 Mb (Figure 1B; Supplemental Table S3), and found assembly size to nearly perfectly mirror genome size estimated from flow cytometry (Pearson’s correlation, r=0.916, p=5.75e-09) (Supplemental Figure S1). The presence of telomere repeats on both terminal ends of chromosome-length contigs in 22 species further supports assembly completeness, as does contig N50 (median 10.2 Mb; range 152 kb-189 Mb) and scaffold N50 (median 72.0 Mb; range 13-198 Mb) (Supplemental Table S2). This contiguity is high in spite of high levels of repetitive transposable element (TE) and tandem repeat sequence, ranging from 54.7-93.3% of assembled sequence (Figure 1B). We generated gene model annotations for each assembly using Helixer (Stiehler et al., 2021), and predicted transcriptomes show a high level of completeness with most (median 98.54, range 95.7%-99.8%) Poales BUSCO genes identified (Supplemental Table S2).

**Figure 1:**
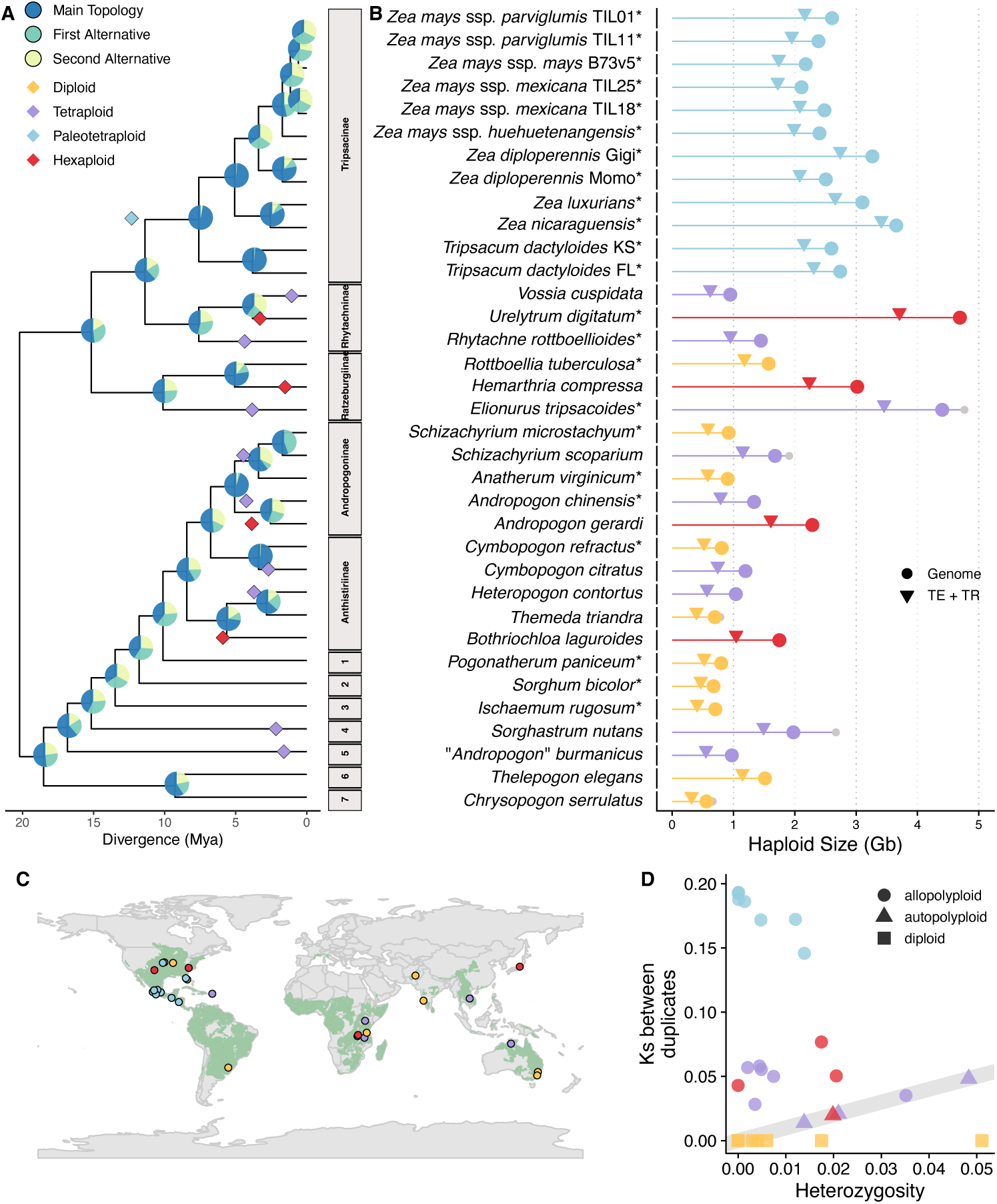
Assemblies of Andropogoneae from throughout the phylogenetic and geographic range have varying divergence, ploidy, and genome size. **A**) Species phylogeny built from 7,725 syntenic genes, including multiple copies in polyploids, using ASTRAL-PRO3. Pie charts at nodes show the quartet support for the main topology in blue, indicated by the tree, the first alternative topology in teal, and the second alternative in yellow. Polyploidy events are shown in diamonds, with colors corresponding to ploidy. Throughout all figures, diploids are shown in yellow, tetraploids in purple, paleotetraploids in blue, and hexaploids in red. The x-axis position of diamonds reflect timing of divergence of parental genomes, so may predate estimated species divergence. The WGD shared by all Tripsacinae is shown with one point at the median parental divergence of all taxa except obligate annual subspecies in *Zea mays*. Subtribes are shown as gray boxes, with names listed when we sampled more than one representative. Single representative subtribes are: 1. Germainiinae, 2. Sorghinae, 3. Ischaeminae, 4. Apludinae, 5&6 are Incertae sedis pending taxonomic revision, and 7. Chrysopogoninae **B**) Haploid size in gigabases of assembly (circle) and TEs and tandem repeats (TR) (triangle), colored by ploidy as in A, with diploids in yellow. Scaffolded assembly size, including N’s, shown with a gray circle. Individuals with * after sample name represent haploid assemblies. Across all assemblies, average assembly size is 1.9 Gb, and average repeat size is 1.5 Gb. **C**) Map showing collection locations for our 33 samples in points colored by ploidy, as in A. The range of all Andropogoneae species is shown in green, constructed from wild occurrences in AuBuchon-Elder et al. (2023). Digitized collections are limited in the Indian subcontinent, although Andropogoneae are abundant there (Welker et al., 2020). **D**) Heterozygosity between alleles versus synonymous substitutions between homeologs for each assembly, with circles designating allopolyploids, triangles autopolyploids, and squares diploids. Diploids, which do not have homeologs, are assigned a Ks value of 0. The gray line indicates a 1:1 relationship between heterozygosity and synonymous substitution rate. *Chrysoposon serrulatus* is excluded from this plot, as it showed elevated nucleotide substitutions arising from nanopore sequencing. As it can be difficult to associate an individual plant with a point, figures with each assembly highlighted are available at https://mcstitzer.github.io/panand_assemblies/.

Combined with publicly available assemblies of *Sorghum bicolor* and *Zea mays* subsp. *mays*, these assemblies offer a dense sample of recent (ca. 0.650 million years of evolution; Tripsacinae (Chen et al., 2022)) and a broad sample of deeper history (ca. 17.5 million years; Andropogoneae (Welker et al., 2020)).

### Relationships between Andropogoneae species

To reconstruct phylogenetic relationships of these genomes, we used gene trees constructed from 7,725 syntenic gene anchors (details provided below) to generate a species tree (Fig 1A). We find pervasive conflict across gene tree topologies, with most bipartitions along the internal branches of the radiation supported by alternative topologies (Figure 1A). This conflict has been noted in previous studies of nuclear and chloroplast markers (Estep et al., 2014; Grass Phylogeny Working Group III, 2024; Welker et al., 2020). The success of these radiations and large effective population sizes likely contribute to these conflicting topologies, as does extensive allopolyploidy in the clade (Estep et al., 2014). Extensive diversification of these species occurred in the late Miocene 12-20 million years ago (Estep et al., 2014) after the origin of C4 photosynthesis (Bianconi et al., 2020). Today they are estimated to cover 15-17% of vegetated land globally (Lehmann et al., 2019).

### Diversity within species reflects differences in reproductive mode and population histories

Comparison of alleles within individuals showed segregating variation (π) across several orders of magnitude (from 0.0002% to 6%; Figure 1D), ranging from nearly homozygous individuals arising from self-fertilization, to high diversity consistent with large effective and census population sizes of these keystone grassland species. For example, low-heterozygosity *Anatherum virginicum* self-seeds extensively, facilitated by cleistogamous flowers (Campbell, 1982), while high-heterozygosity *Heteropogon contortus* is a weedy apomict, found throughout tropical and subtropical latitudes (Carino & Daehler, 1999; Emery & Brown, 1958), and high-heterozygosity *Thelepogon elegans* has populations with permanent translocation heterozygosity facilitated by the formation of ring chromosomes (Sisodia, 1970). In general, heterozygosity values are congruent with expectations from reproductive mode, life history, and range size (Supplemental Text).

### Polyploidy in Andropogoneae is common

We characterized ploidy in our sequenced accessions by counting chromosomes and estimating copy number of syntenic regions derived from whole genome duplications (WGD). Amplification of genomes through polyploidy generates duplicated gene copies, inherited in blocks of conserved syntenic order along chromosomes. The number of times each syntenic region is present in a polyploid can thus give estimates of past WGDs. We used genes from diploid *Paspalum vaginatum* (Sun et al., 2022), which only shares ancestral grass WGDs with Andropogoneae, and is closely related in the sister tribe Paspaleae (Grass Phylogeny Working Group III, 2024). *Paspalum* genes are found in syntenic blocks with 1 to 6 copies in Andropogoneae, after adjusting values based on whether the assembly was haploid or allelic (as described in Li & Durbin, 2024). We integrated our results with cytological literature and chromosome counts to assign ploidy to our samples (Supplemental Figure S2; Supplemental Text). Our studied assemblies include 10 diploids, 9 tetraploids, 4 hexaploids, and 12 paleotetraploids (cytological diploids arising from the Tripsacinae WGD; Figure 1). Their phylogenetic distribution provides fourteen independent polyploid formation events with which to study the impact of polyploidy on genome evolution.

Commonly observed consequences of polyploidy often include reductions in chromosome number, fractionation of duplicate genes, and expansions of transposable elements (Soltis et al., 2016; Wendel, 2015). In our analyses, we see elevated chromosome number in tetraploids and hexaploids (Figure 2A), variable reductions in duplicated genes in all polyploids (Figure 2B), and few expansions of repeat content beyond the multiplication expected from polyploidy (Figure 2C). Given that these patterns deviate from our expectations, we aimed to understand the genomic and temporal factors associated with polyploidy, to understand how these factors influence patterns of genome evolution.

**Figure 2:**
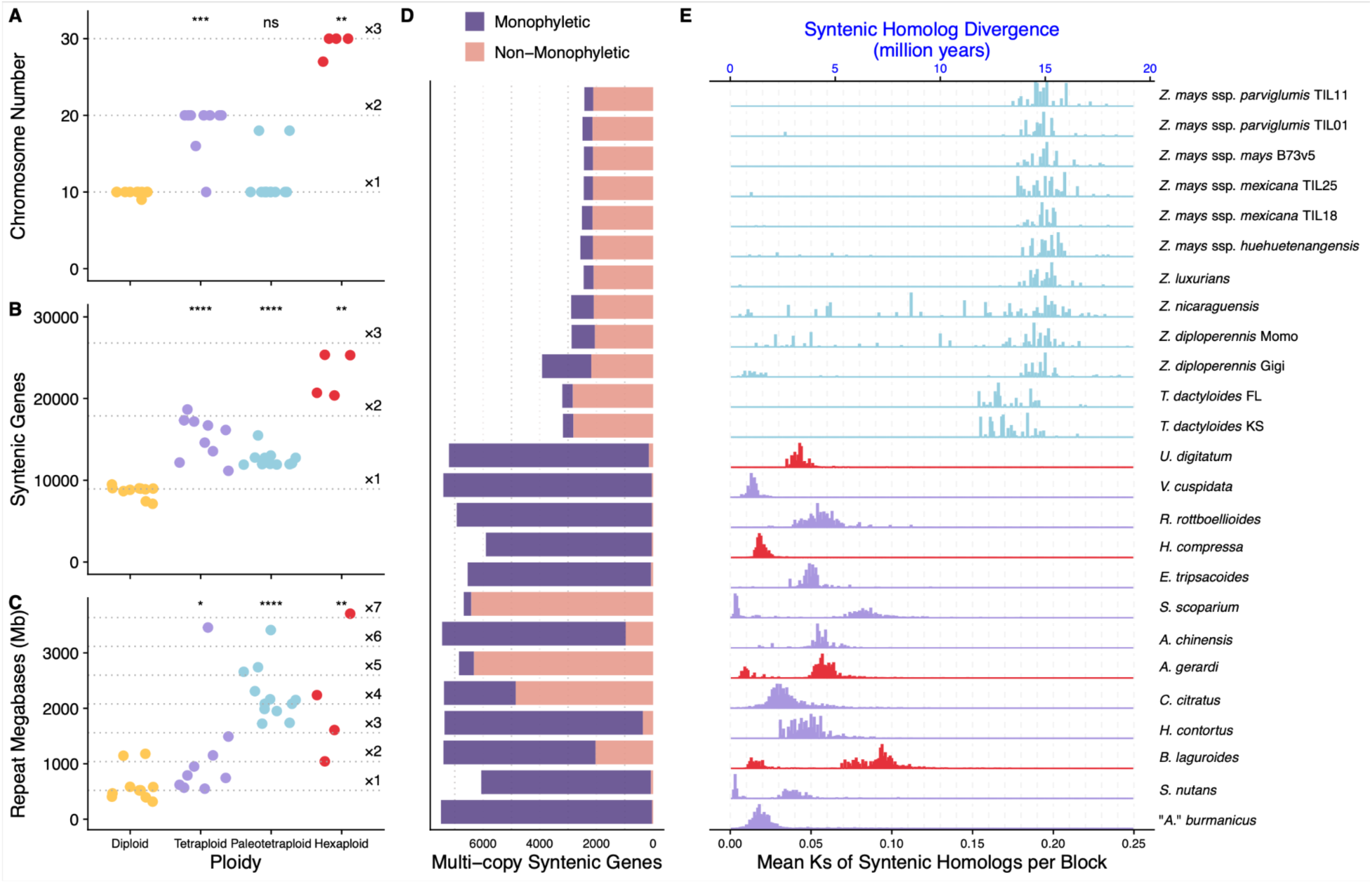
Polyploids in Andropogoneae are abundant. **A)** Haploid chromosome number of each sampled individual, versus ploidy. Diploid median and multipliers to tetraploid and hexaploid expectations are shown with dotted lines. For **A-C**, statistical comparisons of each polyploid group to the diploids were performed using a Wilcoxon rank-sum test. P-values for each comparison shown at top of polyploid group, with *** p<0.001, ** p<0.01, * p<0.05, and ns for nonsignificant. **B)** Number of syntenic genes found in each individual, versus ploidy. Diploid median and multiplier to tetraploid and hexaploid expectations are shown with dotted lines. **C)** Megabases of repeats in each sampled individual, versus ploidy. Diploid median and multipliers up to 7x the diploid median are shown with dotted lines. D) Relatedness of duplicate copies in each polyploid across 7,725 gene trees. Each bar matches labels in **E**. Purple are gene trees where all tips of the given species are monophyletic, and pink are gene trees where the tips are found in paraphyletic or polyphyletic (non-monophyletic) arrangements. As the Tripsacinae paleotetraploidy is shared by multiple species, we downsampled gene trees so only the focal Tripsacinae sample was present in the tree. **E)** Median synonymous substitutions (Ks) between syntenic homologs in polyploids by alignment block, colored by ploidy.

Polyploids form with a mitotic or meiotic “catastrophe” (Comai, 2005), generating gametes with additional sets of chromosomes. Similarity between the subgenomes derived from these parental chromosomes varies from nearly identical (autopolyploid) to fully distinguishable (allopolyploid) (Comai, 2005; Doyle & Egan, 2010; Kellogg, 2016; Ramsey & Schemske, 1998), but this relatedness exists along a continuum. When allelic diversity within subgenomes is indistinguishable from homeologous diversity between subgenomes, the species are likely autopolyploids (Figure 1D; *Vossia cuspidata*, “*Andropogon” burmanicus*, *Hemarthria compressa*, *H. contortus*, *Cymbopogon citratus*; Individual species can be viewed at https://mcstitzer.github.io/panand_assemblies/). Each of these species is mixed-ploidy, and diploid cytotypes are known (Mehra 1982, Darlington and Janaki-Ammal, 1945). Further, *H. compressa* polyploids show formation of multivalents, supporting autopolyploid meiotic pairing (Gupta et al., 2017). Disomic inheritance in polyploids is facilitated by sequence divergence (Bingham, 1980; Mason & Wendel, 2020), as in allopolyploids where allelic diversity within subgenomes is lower than homeologous diversity between subgenomes (Figure 1D). Homeologous diversity in the remainder of Andropogoneae polyploids is consistent with allopolyploidy, with divergent gene copies captured from each parental species (Figure 1D). Allopolyploidy also leaves a signature in gene trees. If the parental taxa are extant and sampled, gene copies in the polyploid will be more closely related to their diploid progenitor than to the other polyploid (homeologous) copy; we call such gene tree patterns “non-monophyletic” with respect to the subgenomes. Our taxon sample places a majority of non-monophyletic relationships of gene copies in four polyploidy events (Figure 2D; Tripsacinae, *Andropogon gerardi*, *Schizachyrium scoparium*, *C. citratus*), strongly supporting allopolyploidy in each case. Two more polyploids have appreciable proportions of non-monophyletic gene trees (Figure 2D; *Andropogon chinensis*, *Bothriochloa laguroides*), likely reflecting our sampling of more distant relatives of potential allopolyploid parents. The distinction between auto and allopolyploidy can be hazy, as evidenced by *C. citratus* with low allelic diversity but with a closely related sampled congener *Cymbopogon refractus*, making a single label difficult.

To estimate the time the parents of these allopolyploids diverged, we used synonymous site divergence (Ks) between homeologous copies (Figure 2E). Using a grass mutation rate (6.5e-9, Gaut et al., 1996) and assuming clock-like substitution rates, median divergences range from 1.05 Mya in *V. cuspidata* (median ks=0.014) to the Tripsacinae WGD at 12.33 Mya (median perennial ks=0.160). Although the Tripsacinae WGD is shared between *Zea* and *Tripsacum* (Estep et al., 2014; McKain et al., 2018), subspecies within *Zea mays* have a derived annual life history (Doebley, 1990; Kempton & Popenoe, 1937), which is expected to increase the number of generations per year and hence estimated divergence (Figure 2E).

The divergence of parental genomes sets an upper limit on the timing of polyploid formation, but does not necessarily reflect when the polyploid nucleus was established (Kellogg, 2016). For instance, while the subgenome progenitors of tetraploid wheat diverged seven million years ago, they formed a tetraploid nucleus only ∼800,000 years ago (Marcussen et al., 2014). Using SubPhaser (Jia et al., 2022), we identified divergent repetitive sequences between homeologous sequences, focusing on subgenome-specific TEs. However, only one assembly, the hexaploid *Urelytrum digitatum*, assigned homeologous sequences into the expected number of groups based on ploidy and chromosome count. In this species, diploid progenitors diverged 1.3 million years before polyploid formation (Supplemental Figure S3). The lack of differentiation in the remaining allopolyploids likely reflects uniform TE invasion and turnover following polyploidization, with insertions distributed evenly across chromosomes.

### Chromosome stability is higher in polyploids than diploids

Chromosome number is a powerful yet simple descriptor of recent polyploidy. Elevated homologous chromosome number poses challenges to cellular processes like mitosis and meiosis (Comai, 2005), so is often countered by chromosome fusions and other rearrangements that reduce chromosome number (Mandáková & Lysak, 2018; Tayalé & Parisod, 2013). Rapid reduction of chromosomes in a newly formed allopolyploid can occur in tens of generations (Buggs et al., 2012; Xiong et al., 2011), and rediploidization via reduced chromosome number appears to be a general consequence of polyploidy, evidenced by its prevalence across flowering plants (Bowers & Paterson, 2021; J. F. Wendel, 2015). Most Andropogoneae genomes, including diploids (Figure 3A), tetraploids (Figure 3B), and hexaploids (Figure 3C) are almost entirely collinear to *Paspalum*, with limited rearrangements. However, paleotetraploid *Tripsacum dactyloides* show more rearrangements (Figure 3D), and all *Zea* individuals (Figure 3D) share a massively rearranged karyotype with a reduction to 10 haploid chromosomes. In contrast to an expectation of rediploidization via chromosome reduction, most Andropogoneae polyploid species show remarkable chromosome stability.

**Figure 3:**
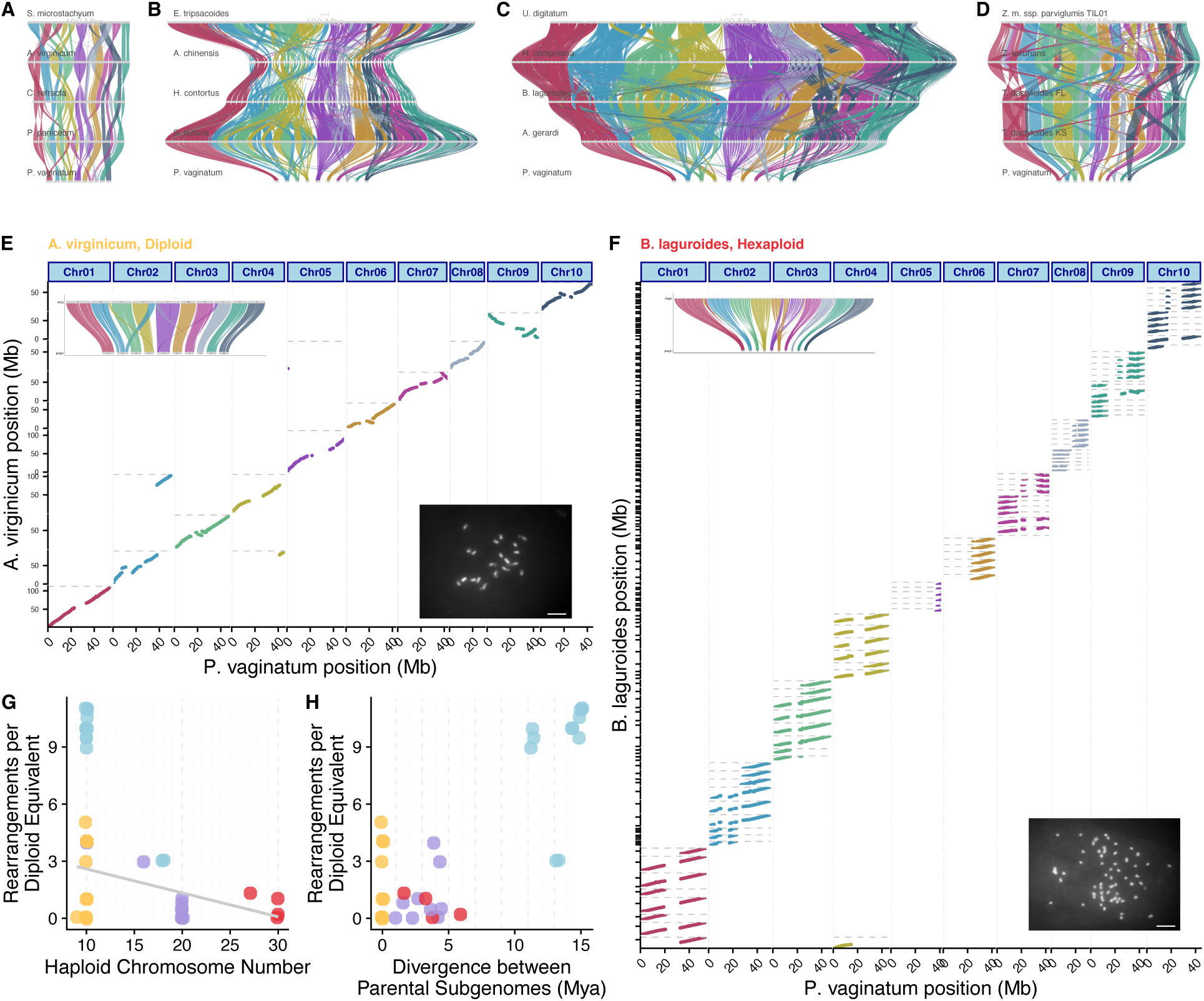
Chromosome stability is higher in polyploids than diploids in Andropogoneae. **A-D** Genespace riparian plots showing synteny of four genomes of each ploidy level, with *Paspalum* on the bottom. Syntenic blocks of each *Paspalum* chromosome are shown as ribbons for **A)** Diploids, **B)** Tetraploids, **C)** Hexaploids, and **D)** Paleotetraploids. **E)** Dotplot of syntenic anchor genes in blocks of 20 or more genes from *Paspalum* chromosomes versus diploid *A. virginicum* chromosomes. The *A. virginicum* assembly is haploid, so only homologous regions are present for each *P. vaginatum* chromosome. Inset in the top left shows a riparian plot comparing chromosomes, and inset in bottom right shows the karyotype of *A. virginicum* with 2n=2x=20, scale bar 10 µm. **F)** Dotplot of syntenic anchor genes in blocks of 20 or more genes from *Paspalum* chromosomes versus hexaploid *B. laguroides* scaffolds. The *B. laguroides* assembly has all six alleles assembled, so each homologous region can be present six times for each *P. vaginatum* chromosome. Inset in top left shows riparian plot comparing assemblies, and inset in bottom right shows karyotype of *B. laguroides* with 2n=6x=60, scale bar 10 µm. **G)** Chromosomal rearrangements show a negative relationship to haploid chromosome number. Each assembly is represented by a point, diploids in yellow, tetraploids in purple, paleotetraploids in blue, and hexaploids in red, as in Figure 1. **H)** Rearrangements are not significantly related to time since divergence of polyploid parents. For **G** and **H**, the median value within each species *Zea* and *Tripsacum* was used for calculating the relationship, due to multiple sampling of this polyploidy.

Contemporary research suggests x=10 is the base chromosome number of Andropogoneae (Spangler et al., 1999), and the majority of polyploids retain multiples of this – only five polyploidy events show chromosome fusions that give rise to reduced chromosome counts (Supplemental Table S3). Among these, only in *Zea* species and *Elionurus tripsacoides* does the reduction reinstate the base diploid chromosome number. To quantify rearrangements, we counted synteny breakpoints that merged two *Paspalum* chromosomes in a single scaffold of each assembly. Diploids have an average of 2 rearrangements, with no significant differences from the mean of tetraploids (1.67 rearrangements), and hexaploids (1.88 rearrangements), while paleotetraploids in *Zea* (mean 20.3 rearrangements) and *Tripsacum* (mean 6 rearrangements) have significantly higher values. However, on a per-chromosome basis, polyploids have fewer rearrangements. When scaled to match a diploid chromosome complement, chromosome number is negatively associated with rearrangement count (R2=-0.416; p=0.035; Figure 3G), suggesting that rather than promoting chromosomal instability, polyploidy stabilizes the chromosome complement. Further, rearrangement abundance does not appear to be solely due to differential amounts of time for rearrangements to occur, as the two do not have a significant relationship (p=0.28; Figure 3H). The underlying mechanisms of this observed retention of elevated chromosome number among most Andropogoneae polyploidy events remain unclear. It may involve selection for gene flow from mixed ploidy populations within the species (Kolář et al., 2017), or a filtering effect that favors the survival of only meiotically and mitotically stable polyploids (Otto, 2007; Ramsey & Schemske, 2002).

The ten *Zea* chromosomes show extensive rearrangements, including several integrations of an entire ancestral chromosome to the center of another (Figure 3D; Supplemental Figure S4), as seen previously (H. Wang & Bennetzen, 2012). Yet aside from inversions and rare translocations, the *Zea* karyotype is unchanged throughout the genus (Braz et al., 2020; Laurie & Bennett, 1985). Of the 14 rearrangements that differentiate *Zea* from *Tripsacum*, seven contain tandemly repeated knob sequences within one megabase of the breakpoint, a major enrichment relative to their average of 1.1% of chromosomal sequence. In maize, chromosomal knobs can act as neocentromeres, mediating meiotic drive by favoring their own segregation (Buckler IV et al., 1999; Dawe et al., 2018; Rhoades, 1942). Such tandem repeats have been linked to genomic shock, as in response to broken chromosomes or cell culture (Lee & Phillips, 1987; McClintock, 1941, 1984; Rhoades & Dempsey, 1972). The tandem array of genes responsible for meiotic drive arose contemporaneously with the divergence of the two genera (L. Chen et al., 2022; Dawe et al., 2018), suggesting meiotic drive may have initiated *Zea*’s pronounced chromosomal rearrangements. However, across Andropogoneae, TE and tandem repeat content is not significantly correlated to rearrangement count (Supplemental Figure S5). These findings suggest that while large tandem arrays are prone to rearrangements, their presence alone does not drive them, as many taxa with large arrays show no rearrangements (Figure 5D). Interestingly, cytological constrictions at knobs have been observed in the genus *Elionurus* (Celarier, 1957; Supplemental Text), the only other instance where we observe a major reduction in chromosome number.

### Most genes are retained in multiple copies, but regulatory sequence turns over rapidly

After a whole genome duplication, functional redundancy leads to decay of homeologous copies, but stoichiometry of protein complexes and pathways leads to preferential retention of some duplicates, particularly transcription factors, developmental genes, and proteins in multi-protein complexes (Birchler & Veitia, 2012; Blanc & Wolfe, 2004; Freeling, 2009). To investigate duplicate retention in our species, we standardized copy number to the genome median, then identified genes showing differential copy number between diploids and polyploids, and conducted a gene ontology (GO) analysis on the top 100 genes. The most strongly enriched GO category included genes encoding ribosomal proteins (“polysomal ribosome,” GO:0042788), consistent with constrained stoichiometry of interacting subunits (Birchler & Veitia, 2012) and with observations of retained duplicates in other plant polyploids (Barakat et al., 2001; Rosado & Raikhel, 2010; Roulin et al., 2013). Signal transduction and stress-response categories were also enriched (Supplementary Table S4). Several developmental processes showed elevated copy number in polyploids, notably root hair initiation, including a gene homologous to brittle culm 10 of rice (Zhou et al., 2009). Increased root hair density following polyploidization has been observed in both *Arabidopsis* (Stetter et al., 2015) and wheat (Han et al., 2016), particularly under nutrient-poor conditions (Salazar-Henao et al., 2016). Other enriched developmental genes relate to cellulose deposition, potentially reflecting the increased mechanical demands on cell walls that accompany altered cell volumes (Corneillie et al., 2019; Morrison, 1980; Serapiglia et al., 2015).

Immediately after polyploidy, all genes exist in multiple copies. Subsequent loss of duplicates can occur through genetic drift (Lynch & Conery, 2000), or may arise from selective processes, such as targeted gene removal (Paterson et al., 2006) or the resolution of dosage conflicts (De Smet et al., 2013; Edger & Pires, 2009). In Andropogoneae, polyploid gene counts suggest such loss, with fewer syntenic genes than suggested from a multiplication of the diploid genome (Figure 2B, 2D, Supplemental Figure S6). To determine whether gene loss is a gradual neutral process, we relate this diploid-equivalent gene count to time since polyploidy, as measured by parental divergence. We observed a negative correlation (p=0.008, R2=0.23, Figure 4E), with a slope implying a loss of 141 genes per million years. However, among tetraploids and hexaploids there is a positive correlation (p=0.03, R2=0.301), such that older polyploids retain more duplicates. This suggests a filtering effect – polyploid lineages that persist over time may do so by maintaining more of their duplicate genes.

**Figure 4:**
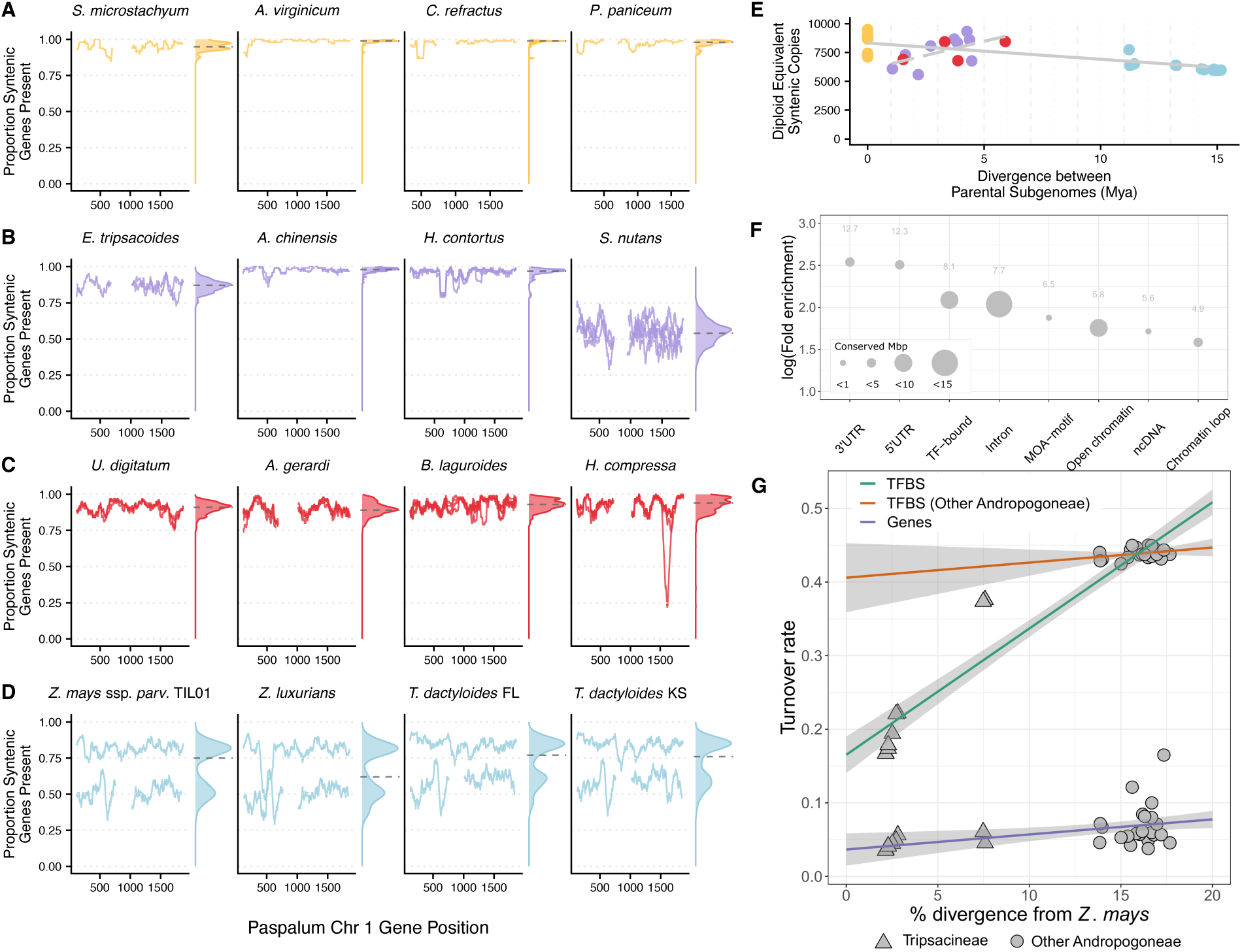
Genes are stagnant while noncoding regulatory sequence turns over rapidly. **A-D)** Retention of syntenic genes in 100 gene windows along *Paspalum* chromosome 1 genes in **A)** diploids with one subgenome, **B)** tetraploids with two subgenomes, **C)** hexaploids with three subgenomes, and **D)** paleotetraploids with two subgenomes. Density plot to the right of each genome shows the genome-wide distribution of these values. **E)** Relationship between gene retention and time since polyploidy, with overall loss (solid line), but a positive relationship within tetraploids and hexaploids (dashed line). Regression for the solid line uses the median value across more densely sampled genera *Zea* and *Tripsacum*. **F)** Enrichment of *Z. mays* genomic features relative to genomic abundance in our non-coding sequence most conserved across Andropogoneae. Absolute fold enrichment is displayed above each point. **G)** Turnover of predicted transcription factor binding sites (TFBS) and genes between *Z. mays* and other Andropogoneae species by genomic divergence. A single representative subgenome was used for TFBS turnover calculations in polyploid species. Loess smooth linear regression lines with 95% confidence intervals are shown. Genomic divergence was calculated using alignments to all fourfold degenerate sites in *Z. mays*.

We next asked whether subgenomic ancestry impacts the gene loss that does occur. When more genes are lost from one homeologous chromosome, it is often attributed to biased fractionation. The maize genome is a classic example of this process, with almost two times as many duplicate genes retained in one subgenome as the other (Schnable et al., 2011). To explore biased fractionation among our polyploids, we measured gene retention in 100 gene windows along *Paspalum* chromosomes for each assembly. Diploids show high retention of syntenic genes (Figure 4A), but most tetraploids (Figure 4B) and hexaploids (Figure 4C) also show near-complete retention of all duplicate copies. In contrast, all Tripsacinae paleotetraploids (Figure 4D) show uneven gene retention across sugenomes: *Tripsacum* retains 85% and 56.8% of genes on each subgenome, while *Zea* retains 81.5% and 49.5%, indicating greater gene loss in *Zea*. Although some polyploids exhibit considerable gene loss (e.g., *S. nutans* retains only 51.4%; Figure 4B), none show the degree of biased fractionation as seen in Tripsacinae. Although lack of biased fractionation may be expected for autopolyploids (Zhao et al., 2017) if recombination equalizes loss across chromosomes, we also found minimal differential fractionation in many allopolyploids. Genome-wide retention of gene copies suggests that polyploidy may act to retain dosage of quantitative traits.

#### Conservation and acceleration of genome-wide conserved elements

Polyploidization duplicates not only genes, but also noncoding regions, expanding the potential for regulatory diversification and expansion (Ebadi et al., 2023; Osborn et al., 2003). To assess the conservation of noncoding sequences, we generated a syntenic multiple sequence alignment of all taxa. We identified 2,302,710 highly conserved Andropogoneae sequence elements covering 72.4 Mbp, enriched in genic regions and potential regulatory sequences associated with binding sites, accessible chromatin, and chromatin loops (Figure 4F, Supplemental Figure S7). After excluding genic sequences (coding sequence, introns, and untranslated regions (UTRs)), we found a set of 1,664,343 conserved non-coding sequences (CNS), each averaging 22 bp, and comprising a total of 36.2 Mbp (Supplemental Table S5), numbers consistent with previous characterizations of CNS in grasses (Liang et al., 2018; Song et al., 2021) and with functional evidence of conserved chromatin accessibility between maize and sorghum (Lu et al., 2019). The Tripsacinae showed substantial sequence-level divergence from other Andropogoneae, with only 34.8% of the non-repetitive maize genome aligning in at least half of the Andropogoneae species sampled, reflecting their deep divergence. Nevertheless, Andropogoneae CNS, including Tripsacinae-accelerated CNS, share features with CNS in other plants including their short size and association with developmental and transcriptional regulation genes (Burgess & Freeling, 2014).

Transcription factor binding sites (TFBS) are an important subset of noncoding sequences and were recently shown to explain the majority of phenotypic variation in many maize traits (Engelhorn et al., 2024). Despite their importance, TFBS turn over more rapidly than genes. For example, in Brassicaceae, CNS turnover is rapid (Haudry et al., 2013) and 74% of experimentally validated TFBS have turned over between *Arabidopsis thaliana* and *A. lyrata* which diverged 10 Mya (Muiño et al., 2016). We investigated whether the Tripsacinae lineage has experienced accelerated evolution in CNS, finding 15,989 CNS with significant signal of acceleration (Supplemental Table S5). GO enrichment analysis revealed that CNS-associated genes were strongly enriched for core physiological and developmental processes (Supplemental Table S6). The Tripsacinae-accelerated CNS were also enriched for terms linked to development including “response to red or far red light” (GO:0009639, fold enrichment=2.7, adjusted p=0.001) and “photoperiodism, flowering” (GO:0048573, fold enrichment=2.1, adjusted p=0.006) (Supplemental Table S7). TFBS found in CNS were most strongly enriched for motifs of the auxin response factor ARF27 (motif MA1691.1, fold enrichment=1.5, adjusted p<1E-300), which is involved in developmental processes (Supplemental Table S8).

To further assess cis-regulatory region turnover across the Andropogoneae, we assessed pairwise TFBS turnover between *Z. mays* and the other Andropogoneae in 11,173 genes meeting our functional and conservation criteria (Methods). TFBS turnover was linked to overall sequence divergence (Figure 4G). Compared to *Z. mays*, the mean pairwise turnover of predicted TFBS is 0.19 in other *Zea* species, 0.37 in *Tripsacum* and 0.44 in other Andropogoneae. In contrast, the coding sequence of these same genes turned over 7 times slower, at a rate of 0.06 across all Andropogoneae. This suggests that the Andropogoneae experience rapid evolutionary turnover of TFBS. Because we focused only on putative regulatory regions that could be aligned between species and thus exclude highly diverged regions, the turnover rates we determined could be underestimates. However, the unexpectedly high rate of turnover of 19% between maize and the closely related teosinte species suggests that a substantial proportion of the predicted TFBS may not be functional, underlining the need for experimentally validated TFBS in maize to improve estimates of TFBS turnover.

#### Genes are further apart in polyploids

This rapid turnover of TFBS over relatively short timescales led us to explore potential drivers, with TEs emerging as key candidates. The maize genome has been described as having tight clusters of genes separated by large swaths of retrotransposons (Fu & Dooner, 2002; Morgante et al., 2005; SanMiguel & Bennetzen, 1998), and across plants, high-density gene regions seem to be evenly spaced independent of genome size (Feuillet & Keller, 1999; Llaca & Messing, 1998). This arrangement suggests that critical regulatory sequences are compressed within short intergenic regions and maintained by selection. We measured median gene-gene distance of syntenic genes across ploidy levels, finding similar values in diploids (4.98 kb) and tetraploids (5.21 kb apart), slightly larger in hexaploids (6.95 kb), but greater in paleotetraploids (19.06 kb) (Supplemental Figure S8). This tighter spacing in diploids and tetraploids aligns with *A. thaliana*’s ∼5kb gene density (Kellogg & Bennetzen, 2004). Gene spacing shows a strong correlation with repeat content (Figure 5A) when excluding paleotetraploids, with genome size alone explaining 72% of the variance in median gene-gene distances (p=2.04e-07). These findings suggest that TE activity may be expanding intergenic regions in paleotetraploid Tripsacinae, introducing new TFBS and disrupting existing ones.

**Figure 5:**
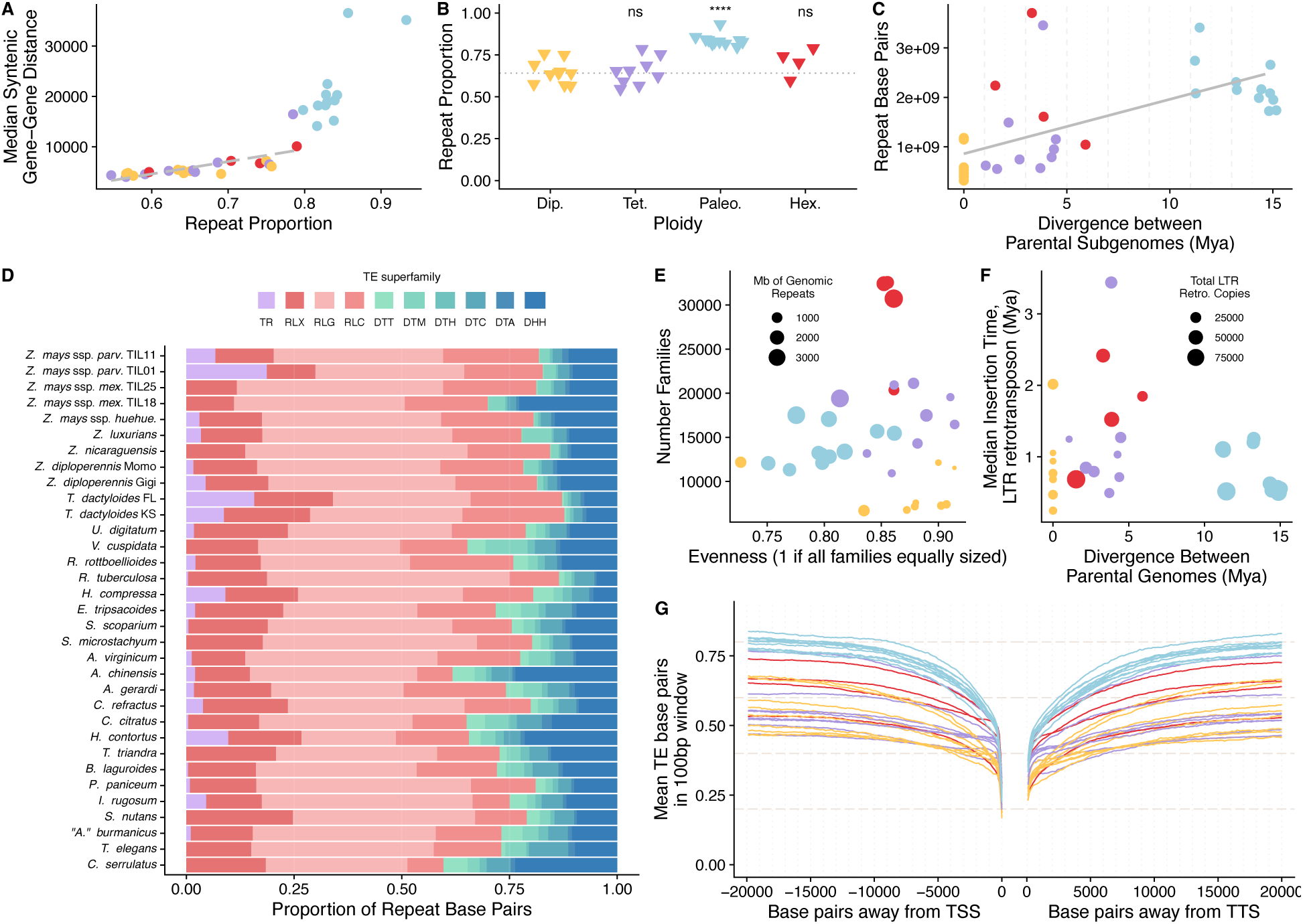
Transposable elements react stochastically to polyploidy. **A)** Proportion of the genome in repeat sequence versus the median distance between syntenic genes. Points are colored by ploidy, with diploids in yellow, tetraploids in purple, paleotetraploids in blue, and hexaploids in red. The dashed line shows regression excluding paleotetraploids, with a strong positive correlation (r= 0.71, p=0.00016). **B)** Repeat proportion related to ploidy level. Statistical comparisons of each polyploid group to the diploids were performed using a Wilcoxon rank-sum test. P-values for each comparison are shown at the top of each polyploid group, with **** p<0.0001 and ns for nonsignificant. Diploid median is shown as a dotted horizontal line. **C)** Divergence between parental subgenomes versus repeat base pairs in each assembly. The solid line includes the median value of *Zea* and *Tripsacum*, which are positively correlated (r= 0.51, p= 0.008). **D)** Proportion of repeats in each genome belonging to different TE superfamilies. TR is Tandem Repeats of all length classes, red colors show LTR retrotransposons (RLX-Unknown; RLC-Ty1/Copia; RLG-Ty3), and blue colors DNA transposons (DTT-Tc1/Mariner; DTM-Mutator; DTH-pIF/Harbinger; DTC-CACTA; DTA-hAT; DHH-Helitron). **E)** Pielou’s evenness metric versus number of families in each assembly. These are calculated only including families with at least 10 copies in the genome. The evenness metric ranges from 0 (one family contributing all copies) to 1 (all families equally sized). Points are scaled by the amount of repeat base pairs in the genome, and colored by ploidy. **F)** Divergence between parental subgenomes and median timing of amplification of LTR retrotransposons. Point size is scaled by the number of structurally intact LTR retrotransposons identified in the genome, and colored by ploidy. **G)** Mean TE base pairs in 100 bp windows away from the transcriptional start site (TSS), left, and transcriptional termination site (TTS), right, of all Helixer genes, colored by ploidy.

### Polyploidy is not associated with TE bursts

Besides polyploidization (Figure 2C), the primary driver of increases in genome size between plant taxa is the amplification of transposable elements (Bennetzen et al., 2005; Bennetzen & Kellogg, 1997; Pulido & Casacuberta, 2023). The genomic shock of allopolyploidy has been hypothesized as a force that can release silencing of TEs, allowing them to expand (McClintock, 1984; Pikaard, 2001). Our data show diploids have lower proportions of their genome coming from TEs (mean=64.8%) than polyploids (mean=75.0%) (p=0.0043) (Figure 5B), but this difference is driven by the paleotetraploid Tripsacinae species and is no longer significantly different after removing them (tetraploid and hexaploid mean=66.9%, p=0.509). This suggests no global expansion of TEs following polyploidy, consistent with certain reconstituted interspecific and intergeneric hybrids (Parisod et al., 2010), young polyploidy events like wheat (Papon et al., 2023), and old polyploidies in animal genomes (Mallik et al., 2023). Despite this general pattern, specific Andropogoneae lineages do exhibit massive TE expansions, most notably *U. digitatum,* which has the largest genome among our samples. Rather than the three-fold expansion expected from hexaploidy, it has over seven times more TE base pairs than expected given the diploid median (Fig 2C). *E. tripsacoides* and *H. compressa* also show large increases in repeat bases relative to diploid multiplications. Thus, while polyploidy may enable TE expansion in some lineages, it does not guarantee it. We modeled TE accumulation as a function of time since polyploidy, and observed a positive correlation, with an estimated gain of 110 Mb of TE sequence per million years (R2=0.23, p=0.008, Figure 5C). This rate exceeds the 38 Mb per million years estimated for diploid rice (Bennetzen et al., 2005; Ma & Bennetzen, 2004), but additional comparative studies across taxa are warranted. Overall, these data do not support global unregulated transposition upon polyploid formation, rather, a gradual accretion of TEs as the polyploid establishes through evolutionary time.

Polyploidy can disrupt the epigenetic environment of the cell (Doyle & Coate, 2019), and these alterations can allow lineages of TEs that were well-silenced in diploid progenitors to exploit novel vulnerabilities in epigenetic silencing. These amplifications are often observed when single TE families reach high copy number (Baduel et al., 2019; J. Chen et al., 2020; Hawkins et al., 2006; Tsukahara et al., 2009). We first identified variation in the proportion of the genome coming from each TE superfamily across Andropogoneae (Figure 5D). As observed in all grass genomes (Vicient et al., 2001), LTR retrotransposons, and particularly Ty3 LTR retrotransposons, contribute the highest proportion of repeat sequence. This simple description of superfamily abundance showed proportional expansions of gene-proximal DNA transposons in specific species, like DTM Mutator elements in *V. cuspidata* and DHH Helitron elements in *Chrysopogon serrulatus* and *A. chinensis*.

To understand whether TE amplifications involved many families or were driven by just a few, we applied species diversity metrics to explore the relative abundance of TE families in each genome. Pielou’s evenness ranges from 0 (where a single family dominates) to 1 (where all families are equally represented). Although evenness varies among genomes (Figure 5E), only paleotetraploids show significantly different evenness compared to other ploidy groups (Wilcoxon rank sum test, W=240, p=2.45e-05). Species with high evenness tend to have lower repeat content (Figure 5E), supporting a lack of successful bursts that overcome epigenetic silencing.

We were curious whether the timing of TE amplification could be linked to polyploidy. However, the rapid turnover of TEs in genomes makes it challenging to directly identify those inserted during polyploidy (Fedoroff, 2012; Vitte & Bennetzen, 2006; Wicker et al., 2018), as repeat landscapes are constantly overwritten. To address this, we estimated the mean age of LTR retrotransposons in the genome by measuring the divergence between their LTRs (SanMiguel et al., 1998). Our analysis reveals a general trend – that diploids have younger LTR retrotransposons, while polyploids harbor older copies, with no clear relationship to the timing of polyploidy (Figure 5F). Young TE insertions have been shown to be more deleterious than older ones (Stitzer et al., 2023), contributing disproportionately to genetic load. As a result, the buffering effect of polyploidy masking deleterious TE insertions may provide an adaptive benefit in these grasses.

Much of the impact of TEs on genome function is via their relationship to generating genetic diversity near genes. We measured the proportion of sequence 20 kilobases upstream and downstream (Figure 5G) of genes that is occupied by TEs. Although all species plateau to values near genome-wide averages 20 kb from genes, the shape of the relationship near genes differs (Figure 5G). For example, diploids and tetraploids reach 50% TEs at a median of 9 kb and 8.4 kb away from the 5’ UTR of the gene, while paleotetraploids (0.5kb) and hexaploids (0.8kb) do so much closer. TEs are well known to affect gene expression (Hirsch & Springer, 2017; Lisch, 2013), disrupting regulatory sequence and introducing new ones. We anticipate incorporating these differences in TE content in key regulatory regions near genes will help understand differences in gene expression.

### Conclusion

Together, these observations underscore a general stasis of genome evolution in these 33 newly assembled Andropogoneae grasses, even under the altered genomic environment of polyploidy. Contrary to the expected patterns of chromosome reduction, gene loss, and TE amplification, most polyploid events deviate from these “rules.” This long-term genomic stability may be facilitated by perennial life cycles of these grasses, which supports persistence on the landscape and exploration of diverse allele combinations, enhancing their adaptive potential.

While polyploidy can sometimes reduce fitness, its broader benefits, beyond simple genome doubling, include fixed heterozygosity, buffering against deleterious genetic load, and increased phenotypic flexibility through gene dosage (Doyle & Coate, 2019; Stebbins, 1971; Tayalé & Parisod, 2013). The large census population sizes, wind-pollination, and high fecundity of Andropogoneae grasses likely enable the exploration of genotypic space, potentially purging genetic load thought to accumulate in polyploids (Haldane, 1933) and exploring gene dosage landscapes.

Modern maize cultivation epitomizes the benefits of hybridization, leveraging heterosis from the combining ability of divergent parental gene pools (Duvick, 2001). Studying how distant relatives of maize adapt to the permanent heterozygosity of polyploidy can provide valuable insights into the complexities of heterosis and its evolutionary significance.

## Methods

### Germplasm

Plant material was sourced from seed banks or collected in the wild (Supplemental Table S1), and grown in the greenhouses at the Donald Danforth Plant Science Center in St. Louis, MO, Cornell University in Ithaca, NY, and the University of California, Davis, in Davis, CA. Permits for collection, export and import were obtained as specified by local governments and nature reserves. Most plants were grown to flowering and voucher specimens were collected and deposited at the herbarium of the Missouri Botanical Garden (MO) and the Australian National Herbarium (CANBR). All specimens were imaged, with the image and metadata uploaded to the Tropicos database following standard protocols (Supplemental Table S1).

### DNA Extraction and Library Preparation

Approximately 5 g of fresh tissue from each plant was extracted for PacBio (Pacific Biosciences, USA) sequencing using one of three high molecular weight DNA approaches. One was based on the Circulomics Big DNA Kit (Circulomics, USA), another on Doyle and Doyle (1987), and another based on the Macherey-Nagel NucleoBond kit (Macherey-Nagel, USA). This DNA was used to generate PacBio libraries, after size selection with either BluePippin (Sage Science, USA), PippinHT (Sage Science, USA), or Ampure beads (Beckman Coulter, USA). Sequencing was completed on the Sequel II or Sequel IIe across 1 to 8 flow cells depending on genome size and output of individual sequencing runs. DNA extraction and sequencing were completed by Corteva Agriscience, Arizona Genomics Institute, and the USDA Genomics and Bioinformatics Research Unit, Stoneville, MS. Nanopore reads for *C. serrulatus* were previously generated (Song et al., 2021).

### Optical Map and HiC Generation

For optical map construction, DNA was extracted from ∼ 0.7 g of fresh leaf tissue from each plant using agarose embedded nuclei and the Bionano Prep Plant Tissue DNA Isolation kit (Bionano, USA). DNA extraction, labeling, and imaging followed the methods previously described in Hufford et al., 2021 and was completed by Corteva Agriscience.

For Hi-C Sequencing, *Tripsacum dactyloides* FL chromatin was crosslinked and isolated from approximately 1g frozen leaf tissue. One Hi-C Seq library was constructed using the Proximo system (Phase Genomics, Seattle) according to the manufacturer’s recommendations and sequenced in a PE150 format in an Illumina Novaseq 6000 analyzer.

### Genome Assembly

We used three sequencing technologies, PacBio Hifi, PacBio CLR, and Oxford Nanopore MinION. Each was assembled into contigs using the detailed methods below (Supplemental Table S2). For 24 samples, we used additional information to further scaffold contigs. This took the form of Bionano optical maps, and HiC for one *Tripsacum dactyloides* individual. For taxa in the genus *Zea*, we used pan-genome anchors to further scaffold into chromosomes.

#### Contig assembly

We generated PacBio Hifi data for 27 individuals, and generated contig assemblies using Hifiasm. HiFi reads obtained from the Circular Consensus Sequencing (CCS) (v6.4.0) pipeline were converted to FASTA format, and HiFiasm (v0.19.5-r590) (Cheng et al., 2021) was used to assemble contigs from the FASTA input reads, using parameters dependent on the scaffolding information available.

For genomes with only PacBio HiFi data, we set the purge level to 3 (-l 3) to generate contigs, purging haplotigs in the most aggressive way. The assembly graph of primary contigs (*.bp.p_ctg.gfa) was used as the representative set of contigs.

For genomes with PacBio HiFi data plus a BioNano optical map, we set the purge level to 0 (-l 0) to prevent the purging of duplicate haplotigs. This generated primary and alternative haplotypes, which were combined to generate contigs.

For the *T. dactyloides* FL genome with HiC data, we used Hi-C partitioning, supplying the HiC FASTQ reads (-h1 -h2), with purge level set to 3 (-l 3). This generated two phased contig assemblies.

We generated PacBio Continuous Long Read (CLR) data for 5 individuals, and generated contig assemblies using Canu (v1.9) (Koren et al., 2017). Pacbio (CLR) data were converted from the native output (Binary Alignment Map, or BAM) format to FASTA format using samtools(v1.17) (Danecek et al., 2021) fasta subcommand, and were then error corrected using Falcon (v1.8.0) (Chin et al., 2016). Briefly, Falcon’s first stage (overlap detection and error correction module) was run, specifying genome size (-genome_size, for auto coverage estimation) and with error correction options of a minimum of two reads, maximum of 200 reads, minimum identity of 70% for error corrections, average read correction rate set to 75%, and maximum seed coverage of 40X, and a chunk size for local alignments of at least 3,000 bp. For the DAligner step, the identical kmer match length was set to 18 bp (-k 18) with a read correction rate of 80% (-e 0.80) and local alignments of at least 1,000 bp (-l 1000). Contigs were generated using Canu (v1.9), after merging the error-corrected reads from Falcon jobs, using the default options except for ovlMerThreshold=500 (kmers that occur more than 500 times are not used as seeds).

We used Oxford Nanopore MinION reads for one individual, *Chrysopogon serrulatus* (Song et al., 2021). Basecalling was performed using Guppy (v 2.1.3), and FASTQ files were used for error correction. The porechop package (Wick et al., 2017) was used to clean for adapter trimming and error correction of ∼ 52 Gb of MinION reads. The reads were then assembled using Canu v1.8 with the default parameters as described above in the section on CLR, but with the default ovlMerThreshold.

#### Scaffolding

We generated Bionano optical maps for 21 individuals. The Bionano optical maps were processed using Bionano Solve (v3.4) and Bionano Access (v1.3.0), following the methodology outlined in Hufford et al., (2021). For the hybrid assembly, default settings from the configuration file (hybridScaffold_DLE1_config.xml) and parameters file (optAr-guments_nonhaplotype_noES_noCut_DLE1_saphyr.xml) were utilized. The scaffolding phase in Bionano Solve incorporates 1) estimated gaps of varying N-size, excluding 100 bp or 13 bp gaps, as determined through calibrated distance conversion of the optical map to base pairs, 2) unknown gaps (100-N gaps), and 3) 13-N gaps, which are introduced when two contigs overlap. Due to polyploidy and high heterozygosity of many genomes, the 13-N gaps were curated manually, using Bionano Access (v1.3.0). Alignments of contigs to the optical map were examined in detail, and contigs were either trimmed near overlapping regions or exact duplicates were labeled as alternative haplotypes (e.g. alt-scaf_NNN).

We generated Hi-C reads from *Tripsacum dactyloides* FL. Hi-C reads in FASTQ format were first mapped to both haplotypes of the phased haploid genomes of *Tripsacum dactyloides* FL (contigs assembled using Hi-C partitioning) using the Burrows-Wheeler Aligner (BWA) (v0.7.12) (H. Li & Durbin, 2009). The juicer pipeline (v1.6) (Durand et al., 2016) was used to filter out erroneous mappings (MAPQ = 0) and duplicates, and to generate the interaction matrix. The 3D-DNA pipeline (v180) (Dudchenko et al., 2018) was then used to anchor the contigs to chromosomes and error correct the contigs using default parameters. The resulting Hi-C contact maps were manually examined using JUICEBOX Assembly Tools (v2.15.07) (Dudchenko et al., 2018) and a few out-of-place contigs were manually corrected. A final assembly for each phased haplotype was generated and the genome with fewer Ns was designated as primary haplotype.

To scaffold the *Tripsacum dactyloides* KS genome, we used ALLMAPS (Tang et al., 2015) to order and orient the primary and alternative contigs. Briefly, haplotype 1 of *T. dactyloides* FL was aligned against *T. dactyloides* KS. Randomly sampled regions of the alignment were used as markers for input into ALLMAPS.

For nine individuals in the genus *Zea*, pan-genome anchor markers were used to further scaffold contigs or scaffolds to chromosomes, as in Hufford et al. (2021).

### Genome size estimation

We estimated genome size of sequenced individuals outside the Tripsacinae using methods modified from (Doležel et al., 2007), and described in Phillips et al., (2023). Two internal standards were used, depending on the reference: maize B73 inbred line (5.16 pg/2C) and our *A. virginicum* accession (2.17 pg/2C). We placed approximately 10 × 1 cm of fresh leaf tissue for the target and sample standard in a plastic square petri dish, and added 1.25 mL of a chopping solution composed of 1 mL LB01 buffer solution, 250 *μ*L propidium iodide (PI) stock (2 mg/mL), and 25 *μ*L RNase (1 mg/mL) (Doležel et al., 2007). We next chopped the tissue into 2–4 mm lengths and mixed the chopping solution through the leaves by pipetting. The solution was then pipetted through a 30 *μ*m sterile single-pack CellTrics filter into a 2 mL Rohren tube on ice. At least three replicates were chopped separately and analyzed for each individual. The samples were left to chill for 20 min before analysis with a BD Accuri C6 flow cytometer. Samples were run in Auto Collect mode with a 5-minute run limit, slow fluidics option, a forward scatter height (FSC-H) threshold with less than 200,000 events, and a one-cycle wash. The cell count, coefficient of variation of FL2-A, and mean FL2-A were recorded for the target and reference sample with no gating. Results were analyzed separately for each replicate and manually annotated to designate the set of events. We averaged values across all replicates of each individual (Supplemental Table S3).

### Chromosome counts

Newly formed root tips were harvested from greenhouse-grown reference plants approximately one week after transplanting to new growth medium. Two or more roots were examined for each species. Exposure to nitrous oxide (160 psi for 2.5 - 3 hr) was used to stop mitosis in metaphase (Kato, 1999). Methods used for fixation, enzymatic digestion of meristematic tissue, and slide preparation have been described in detail (Kato et al., 2011; Phillips et al., 2023). The digestion times and amount of acetic acid-methanol solution used to resuspend the digested meristem varied based on meristem size. The cross-linked suspension was stained with a 1/20 dilution of Vectashield with 4’,6-diamidino-2-phenylindole (DAPI) (Vector Laboratories, Burlingame, CA). Images were acquired with Applied Spectral Imaging (ASI) software (Carlsbad, CA) on an Olympus BX61 fluorescence microscope and saved in grayscale. The background was reduced using Adobe Photoshop Brightness/Contrast and/or Curves functions. Some images were sharpened with either ASI or Microsoft PowerPoint software.

### Gene annotation and homolog identification

We used Helixer (Stiehler et al., 2021), a deep learning gene prediction model to produce annotations for each genome. We used the plant model, trained on 51 land plant genomes. As these models often generate false positive gene annotations, often including transposable elements in the gene set, we aimed to filter these annotations. We ran Orthofinder v2.5.5 (Emms & Kelly, 2019) within GENESPACE v1.3.1 (Lovell et al., 2022) on all assemblies and the *Paspalum vaginatum* outgroup. We retained orthogroups with >40 and <200 copies, generating an “orthology filtered gene set,” which we use for analyses involving gene copy number (detailed in *Gene Ontology searches* section).

We generated another set of “traditional” gene annotations using *ab initio* predictions from BRAKER (v2.1.6) (Brůna et al., 2021), direct evidence inferred from transcript assemblies using the BIND strategy (Li et al., 2022), and homology predictions using *Sorghum bicolor* and *Zea mays* subsp. *mays* (B73v5) annotations were generated using GeMoMa (v1.8) (Keilwagen et al., 2018) Annotation Filter tool. Predictions were prioritized using weights, with the highest for homology (1.0), followed by direct evidence (0.9), and the lowest for gene predictions from *ab initio* methods (0.1). Weights were assigned based on reliability, and the Annotation Filter ensured prediction completeness, external evidence support, and RNAseq support. The canonical transcript for each gene was predicted using TRaCE (Olson & Ware, 2021).

Traditional gene annotation pipelines can struggle in polyploid and allelic assemblies, due to multiple mapping across homologous copies. Additionally, the close gene spacing in some of our assemblies led to inappropriate merging of adjacent gene models, particularly in genomes less than 1 gigabase. We provide these traditional gene models for consistency with community standards, but for comparisons across taxa and analyses in this paper, we use the “orthology filtered gene set” derived from Helixer models and a set of syntenic anchor genes described below in analyses throughout this paper.

### Repeat annotation

We ran EDTA (Ou et al., 2019) with default parameters on each assembly, generating a repeat library and gff annotation. We used the “traditional” gene annotations as input to EDTA. As no gene annotation was produced for *Zea luxurians*, we supplied the B73v5 gene sequences for masking purposes. Although EDTA can be supplied with a reference TE library, such curated libraries are only available for maize. In order to compare TEs across our taxonomic sampling, we performed de novo searches on each assembly, so methods were consistent.

To calculate TE family evenness, we used Pielou’s evenness metric (Pielou, 1966), counting the number of copies in each TE family in each genome. Pielou’s evenness is a ratio between the Shannon Index and the hypothetical value if all families had the same relative abundance.

### Tandem repeat identification and masking

Initial investigation of EDTA outputs suggested many megabases of *Zea* knob sequences were annotated as different TE superfamilies in each assembly. To identify tandem repeats that may be falsely annotated as TEs, we used TRASH (Wlodzimierz et al., 2023) with default parameters to annotate tandem repeats in each assembly. Within each assembly, we filtered to unique primary consensus sequences at least 40 bp long and found in at least five positions (discontinuous tandem arrays). We used these sequences as a repeat library to mask the assembly it was generated from, using RepeatMasker v.4.1.0 (Smit et al., 2013) with parameters (-q -no_is -norna -nolow -div 40), generating gff output with (-gff). We merged this RepeatMasker output with the EDTA gff, first using bedtools subtract to remove EDTA TEs that overlapped tandem repeats, then concatenating this output with the tandem repeats. There was one remaining knob-related sequence incorporated into a Ty3 family that makes up ∼200 Mb of sequence in *Z. nicaraguensis* (TE_00015576), which we removed for analyses of TE content.

To identify terminal telomere sequence, we searched contigs greater than 1 Mb for the Poales AAACCCT telomere repeat using tidk (Brown et al., 2023). We consider the telomere present at a sequence end if >100 repeats are present in the terminal 30 kilobases.

### Gene synteny identification

We used AnchorWave (Song et al., 2022) to generate syntenic paths through each assembly, with values informed by evidence of polyploidy (Supplemental Table 3). We used the haploid assembly and gene annotation of the non-Andropogoneae outgroup *Paspalum vaginatum* (v3.1; Sun et al., 2022) as the reference, and allowed each gene anchor to participate in up to 2 paths for diploids, 4 for tetraploids and paleotetraploids, and up to 6 for hexaploids. In cases of uncertain ploidy, we increased the number of paths allowed to 6. For each *Paspalum* gene, we counted how many paths it participated in per taxon, and recorded the start/end coordinates of each genic CDS alignment. We filtered to anchors present in at least one copy in 32/35 assemblies (>90%), to generate a set of 9,168 conserved syntenic anchors, which we refer to as our “syntenic gene anchors.”

Due to the extensive chromosome collinearity between *Paspalum* and Andropogoneae, we used these syntenic blocks to count chromosome rearrangements in each assembly. We used syntenic blocks containing at least 30 genes, and considered each contig that contained syntenic blocks that matched two *Paspalum* chromosomes as a rearrangement. We excluded four low-contiguity assemblies from these calculations (*Thelepogon* elegans, *“Andropogon” burmanicus, Rhytachne rottboelloides*, *Cymbopogon* citratus).

### Ploidy

To estimate ploidy, we integrated cytology, gene synteny counts, heterozygosity, and literature reports. Historical conflict over the base chromosome number of Andropogoneae can complicate interpretations, as n=5 and n=10 can be indistinguishable when conflated with different ploidy levels. For example, a n=5 tetraploid would have 20 pachytene chromosomes, as would a n=10 diploid. Additionally, whether alleles are assembled into distinct contigs impacts the depth of syntenic blocks, as a tetraploid with alleles collapsed into two contigs would have a depth of 2, just as a diploid with both alleles assembled would. Using literature searches of chromosome counts and our own chromosome squashes as a guide to possible ploidy, we first assigned assemblies as allelic tetraploids or hexaploids if their synteny depth reached 4 or 6, and haploid hexaploids when the modal synteny depth was 3. As we measured 1C genome size, an allelic assembly will have ∼2 times the Mb of DNA as the flow cytometry measurement, allowing us to further classify tetraploid and diploid taxa. Our final assignments of ploidy are shown in Supplementary Table S3, and haploid assemblies are designated with a (*) after their name in Figure 1. When presenting genome assembly size in Figure 1B, we divide allelic assemblies by two to present haploid size, and when presenting gene counts and repeat content throughout the paper, we divide values for allelic assemblies by two to present comparable haploid equivalents.

### Heterozygosity

For taxa with haploid assemblies, we mapped raw reads back to the reference genome using minimap2 (H. Li, 2018) with -ax map-pb, and called SNPs using DeepVariant 1.6.1 (Poplin et al., 2018). We used the number of heterozygous SNPs in the resulting gvcf divided by homozygous reference calls in the same gvcf as a measurement of heterozygosity. This helped classify ploidy for two taxa with ambiguous assignment, *E. tripsacoides* and *R. tuberculosa*. As both these assemblies showed very low heterozygosity, we assigned *E. tripsacoides* as a haploid assembly of a tetraploid, and *R. tuberculosa* as a haploid assembly of a diploid. However, such a result could arise after many generations of selfing, which may be possible for congener *Rottboellia exalta* (Supplemental Text). We do not report heterozygosity values for *C. serrulatus*, as it appears to have elevated heterozygosity estimates arising from base errors of early generation Nanopore reads.

### Gene tree reconstruction

For each *Paspalum* syntenic anchor, we extracted the corresponding genomic sequence from each assembly using samtools faidx (Danecek et al., 2021). From these syntenic anchors, we combined each sequence into a multiple-fasta of CDS and intronic sequence of each anchor gene, and aligned them with MAFFT (parameters --genafpair --maxiterate 1000 -- adjustdirection) (Katoh & Standley, 2013). We generated a gene tree from this multiple sequence alignment using RAxML (Stamatakis, 2014) with 100 rapid bootstrap replicates (parameters -m GTRGAMMA -p 12345 -x 12345 -# 100 -f a). As we included introns, some alignments failed to align due to memory issues, so we produced gene trees for 7,725 syntenic anchors. We note that this set of genes is not reliant on the gene annotation of each assembly, so it likely captures both genes and pseudogenes.

To calculate synonymous diversity between copies, we extracted codon positions based on the alignment and the *Paspalum* gene used as an anchor. We calculated all pairwise comparisons of all tips in the gene tree and filtered to gene copies within a polyploid to characterize intraspecific Ks values.

### Species tree reconstruction

We adjusted the tip labels in each gene tree to be the species name, such that gene trees were multilabelled for allelic and homeologous copies, and provided these 7,725 gene trees as input to ASTRAL-PRO3 (Tabatabaee et al., 2023; Zhang et al., 2020) using default parameters.

### Gene Ontology searches

We used Blast2GO (Conesa et al., 2005) to generate Gene Ontology categories for each Helixer gene in each species, then merged GO terms across all copies within an orthogroup. We tested for enrichment using TopGO (Alexa & Rahnenführer, 2009). We identified a set of orthogroups with copy number deviations from the standardized genome count for polyploids vs diploids. To determine whether gene copy number deviated between the groups, we first calculated the median copy number for each gene within each assembly. We then standardized the assembly’s copy number by subtracting this median. For each orthogroup, we used a Wilcoxon rank-sum test to compare deviations in copy number between the groups. P-values were obtained for each orthogroup-specific comparison, and ranked to select the top 100 genes for exploration with GO via TopGO.

### Subgenome phasing

We ran SubPhaser (Jia et al., 2022), with a k-mer length of 17 and a minimum k-mer count of 200. Homologous sequences were defined based on the AnchorWave syntenic regions.

### Conservation and acceleration of genomic elements at different phylogenetic scales and ploidies

Andropogoneae genomes soft-masked for EDTA repeats were aligned with Cactus 2.1.1 (Armstrong et al., 2020) and a multiple alignment based on the B73 reference was extracted from the alignment graph. Syntenic alignments to B73 were retained based on MCScanX (Y. Wang et al., 2012) syntenic gene blocks using traditional gene annotations. No gene annotations were generated to annotate the *Zea luxurians* genome, so the annotation used for MCScanX was generated by lifting over the *Z. nicaraguensis* annotation using LiftOff 1.6.2 (Shumate & Salzberg, 2021) with the “polish” flag. Chimeric subgenomes were assigned for all polyploid species based on MCScanX synteny to the ancestral *Paspalum vaginatum* genome, clustering scaffolds syntenic to each *P. vaginatum* chromosome into subgenomes based on a custom greedy algorithm that minimized overlap between subgenomes. The resulting subgenomes are chimeric as each chromosome may be composed of different biological subgenomes. We do not expect this to impact our analysis of conserved elements as divergence between subgenomes was relatively low.

A neutral model of evolution was fit to the Andropogoneae phylogeny using fourfold degenerate sites from maize chromosome 10 with phyloFit from the PHAST 1.4 package (Hubisz et al., 2011). A set of most conserved elements was generated using the PhastCons “most-conserved” flag from the PHAST package with an expected length of 8bp, after training to generate models of conserved and non-conserved elements using genome-wide multiple alignments with “--coverage 0.45”. To prevent reference-bias in the discovery of CNS, the B73 reference was masked and all other Tripsacinae were excluded for the phastCons analyses. The resulting 2,302,710 conserved elements were then filtered to exclude elements shorter than 5 bp or with >= 1 bp overlap with CDS, introns, or untranslated regions (UTR). In addition, conserved elements with BLASTX hits with e-value <= 0.01 to the Swissprot Viridiplantae protein database were removed, to ensure unannotated genes and pseudogenes were excluded.

To determine the presence of a CNS element in each Andropogoneae genome, we required an alignment covering at least 50% of the element, excluding gaps. CNS were classified as “downstream”, “upstream”, “downstream distal”, or “upstream distal” based on the B73 gene annotation and a threshold of 1 kbp distance from the nearest gene feature to determine whether a CNS was distal. Fold enrichment of genomic features in the conserved elements was calculated following Song et al., (2021) by dividing the proportion of bp of a feature that were conserved by the proportion of bp in the genome that overlap the feature. Chromatin loops were determined based on HiC data (Ricci et al., 2019), and accessible chromatin regions (ACR) were based on single-cell ATAC experiments from (Marand et al., 2021). MNase-defined cistrome-Occupancy Analysis motifs were obtained from Savadel et al., (2021) and transcription factor bound regions were based on ChIP-seq provided by Tu et al., (2020). Other genomic annotations including of TEs and noncoding RNA were based on the MaizeGDB B73v5 genome annotation. We used phyloP with “--method LRT --mode ACC” from the PHAST package to test for lineage-specific acceleration in Tripsacinae in elements conserved across Andropogoneae. Multiple testing correction of LRT p-values was conducted using the Benjamini-Hochberg method with an FDR threshold of 0.05. A GO enrichment analysis of all CNS and Tripsacinae-accelerated CNS was conducted using rGREAT 1.1.0 (Gu & Hübschmann, 2023) using the B73 v5 annotation and default parameters. For Tripsacinae-accelerated CNS the CNS was used as the background. GO terms with a fold enrichment <2 and an adjusted p-value >=0.1 were filtered.

We assessed the turnover of TFBS across Andropogoneae based on alignments to B73 predicted TFBS in the multiple alignment as well as predicted TFBS for each genome. TFBS were predicted based on the 46 representative plant motifs in the JASPAR 2022 plant-specific database (Castro-Mondragon et al., 2022), which were trimmed using universalmotif 4.3 (Tremblay, 2024) with a minimum allowed information content of 0.5 bits. Motif scanning was conducted using a custom kotlin script with a detection threshold of 70% of the maximum position weight matrix score. To focus on the potentially most functionally relevant TFBS, we used TFBS in the 1kb region upstream of the translation start sites of B73 genes, which also met the following criteria: has gene expression >0 TPM (Hufford et al., 2021), is a core gene across maize NAM lines (Hufford et al., 2021), is not a tandem duplicate in B73, the 1kb upstream of the translation start site intersects with >=1 ChIP sequencing peak from data generated by Tu et al. (2020), and has >=1 syntenic collinear ortholog across the other Andropogoneae species. For each Andropogoneae query species, we compared the TFBS in the 1kb region upstream of the translation start sites of the selected B73 genes and the TFBS in the 1kb region upstream of the translation start sites of the collinear gene in the query species, selecting a single random ortholog if there were multiple orthologous collinear genes in the query species. A single representative subgenome for each polyploid query species was used. To account for limitations of the alignment and structural variation, a reference TFBS was considered present in a query species if it was aligned and a matching prediction was present or if it was not aligned but a matching prediction was present in the query region. To compare the TFBS turnover, genic turnover was also calculated using the Orthofinder orthogroups for the same set of genes analyzed for TFBS turnover, excluding orthogroups with multi-copy B73 genes. *Z*. *nicaraguensis* was excluded from these analyses because its gene annotation was incomplete due to missing sequences in the main haplotype assembly, and *Z*. *luxurians* was excluded because it did not have a high-quality gene annotation. To compare the turnover rates with the phylogenetic distance to *Z*. *mays*, sequence divergence to each species was calculated using the set of fourfold degenerate sites used to calculate the neutral model.

## Data Availability

Raw data will be available under NCBI/EBI BioProject PRJEB50280 and genome assemblies at USDA Ag Data Commons upon publication. The code used to generate assemblies and conserved noncoding sequence analyses is available at https://github.com/HuffordLab/panand_genome_evolution, and code for other analyses and figures at https://github.com/mcstitzer/panand_assemblies. Additionally, figures highlighting each plant can be viewed at https://mcstitzer.github.io/panand_assemblies/

## Acknowledgements

This article is based upon work supported by the National Science Foundation under Grant Number 1822330, and the U.S. Department of Agriculture–Agricultural Research Service (USDA-ARS) under Project Number 5030-21000-072-00-D (Corn Insects and Crop Genetics Research Unit, Ames, Iowa), Project Number 6066-21310-005-00-D (Genomics and Bioinformatics Research Unit, Stoneville, MS), and Project Number 8062-21000-043-00-D (Plant, Soil and Nutrition Research Unit, Ithaca, NY). Mention of trade names or commercial products in this publication is solely for the purpose of providing specific information and does not imply recommendation or endorsement by the U.S. Department of Agriculture. USDA is an equal opportunity provider and employer.

A.S. and A.C.S were supported by the National Institute of General Medical Sciences, National Institutes of Health, under award number R01GM102192. M.C.S. was supported by the National Science Foundation Postdoctoral Research Fellowship in Biology under Grant Number 1907343.

This research used resources provided by the SCINet project and the AI Center of Excellence of the USDA Agricultural Research Service, ARS project numbers 0201-88888-003-000D and 0201-88888-002-000D and The Texas Advanced Computing Center (TACC) at The University of Texas at Austin. Computational resources and data management were provided by the Bioinformatics Facility (RRID:SCR_021757) at the Cornell Institute of Biotechnology. The authors acknowledge financial support from Inari Agriculture for the sequencing of six species included in this project, and financial support from Corteva Agriscience for sequencing of ten individuals.

Lynn Clark (Iowa State University) provided germplasm for *Pogonatherum paniceum*. We acknowledge all who have contributed to the conservation, cultivation, and study of the natural diversity of Andropogoneae grasses.

## Notes

https://mcstitzer.github.io/panand_assemblies/

### Competing Interest Statement

The authors have declared no competing interest.

