## Supplemental Figures for "Extensive genome evolution distinguishes maize within a stable tribe of grasses"

**Supplemental Figure S1. Assembly size is correlated with flow cytometry estimates of genome size.** Relationship between haploid assembly size and genome size as estimated for flow cytometry. We did not calculate flow cytometry estimates for paleotetraploid Tripsacinae.

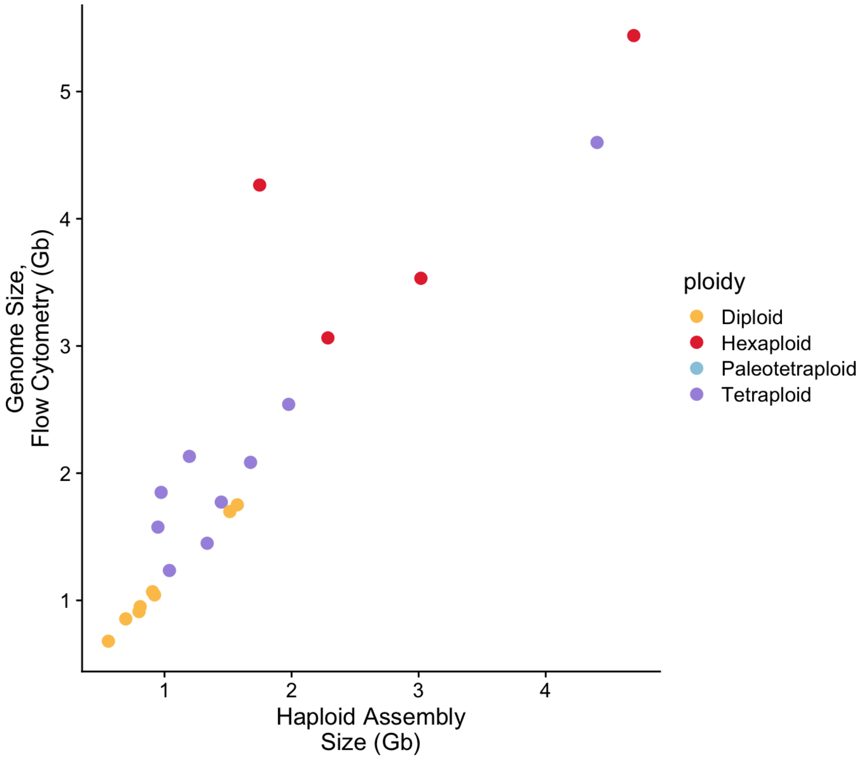

**Supplemental Figure S2. Metaphase chromosomes from the root tips of nine grass species listed alphabetically**. **a**, *Andropogon gerardii*, 2n = 60. **b**, *Andropogon virginicus* 2n = 20. **c**, *Bothriochloa laguroides*, 2n = 60. **d**, *Cymbopogon refractus*, 2n = 20. **e**, *Elionurus tripsacoides* 2n = 20. **f**, *Heteropogon contortus*, 2n = 40. **g**, *Schizachyrium scoparium*, 2n = 40. **h**, *Sorghastrum nutans*, 2n = 40. **i**, *Themeda triandra*, 2n = 20. Scale bars, 10 µm.

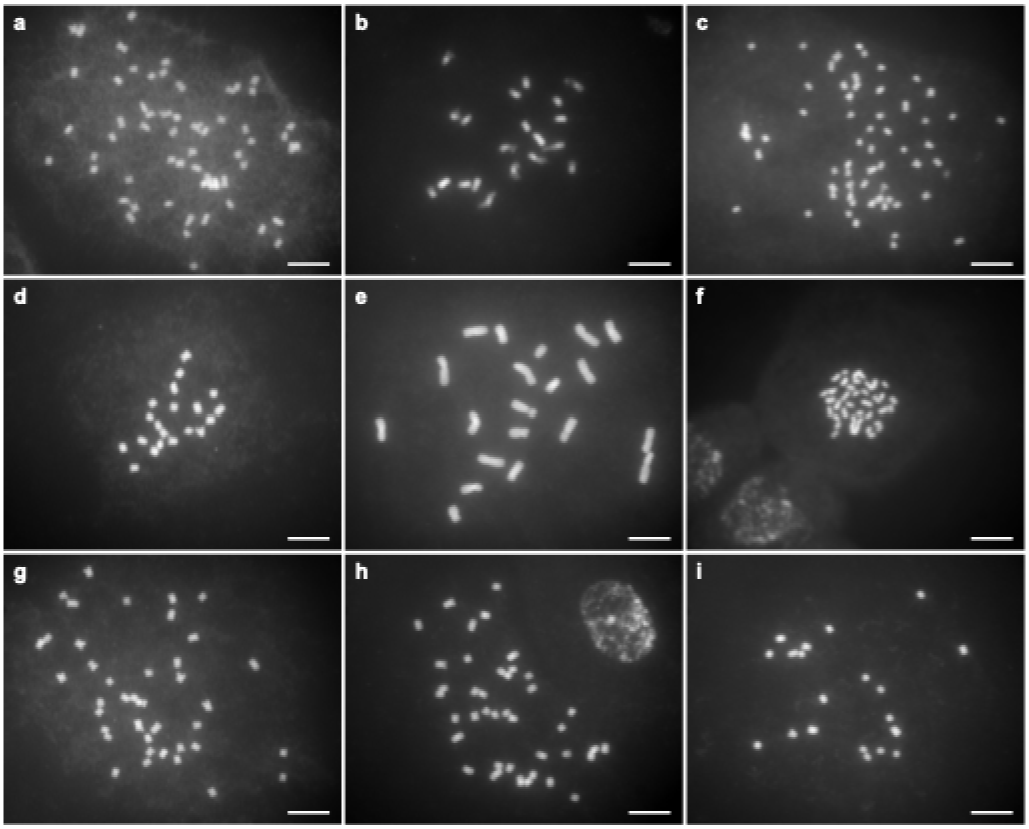

**Supplemental Figure S3. Distributions of timing of insertion of subgenome-specific TEs in *U. digitatum*.** Subgenome specific 17-mers were used to define each subgenome using SubPhaser, then identify subgenome-specific LTR retrotransposons.

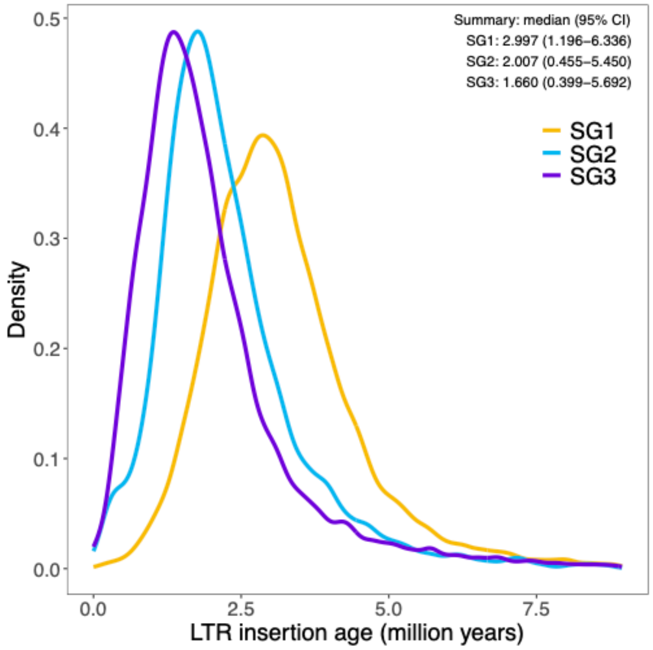

**Supplemental Figure S4. Dotplot of chromosomal continuity between *Zea* and *Tripsacum*.** Zea mays subsp. *mays* B73 on the x-axis, with syntenic positions shared with *Tripsacum dactyloides* FL-hap1 on the y-axis. Lines colored by *Paspalum* chromosome, as shown in Figure 3.

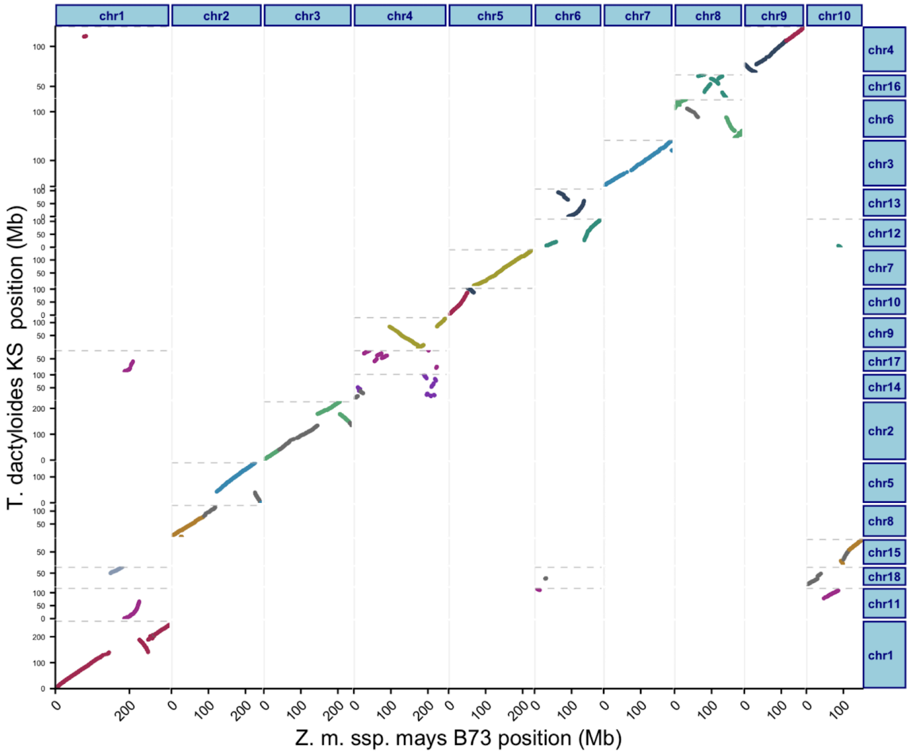


**Supplemental Figure S5. Rearrangements are not correlated with repeat content**

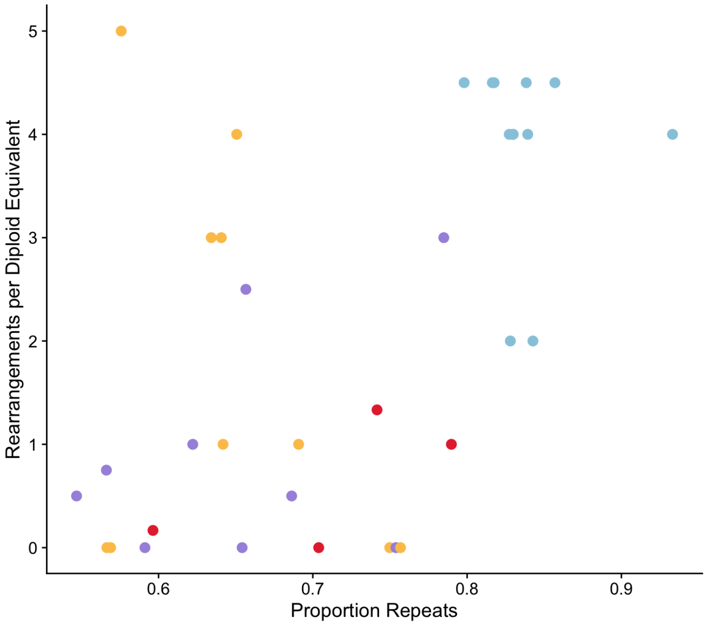

**Supplemental Figure S6. Helixer and orthology-filtered gene counts. A)** Copy number of all Helixer genes in a haploid assembly. **B)** Orthology filtered Helixer genes, from orthogroups with >40 or <200 gene copies.

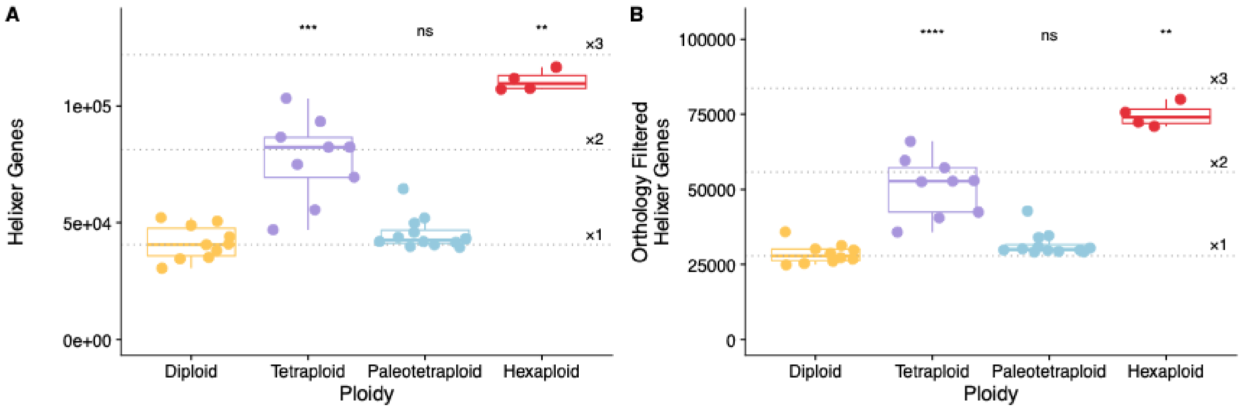

**Supplemental Figure S7. CNS and alignment validation A)** A total of 33 species were aligned to the *Z. mays* reference genome (2131.8 Mbp in total length, of which 271.7 Mbp is not composed of transposable elements). **B)** Metaplot of distribution of Andropogoneae conserved non-coding sequence (CNS) distances from the nearest gene. Distances for CNS downstream of genes are shown as negative values. **C)** Andropogoneae phylogeny with CNS content for each species. Distal sequences are defined as those >1 kbp distant from the nearest *Z. mays* gene.

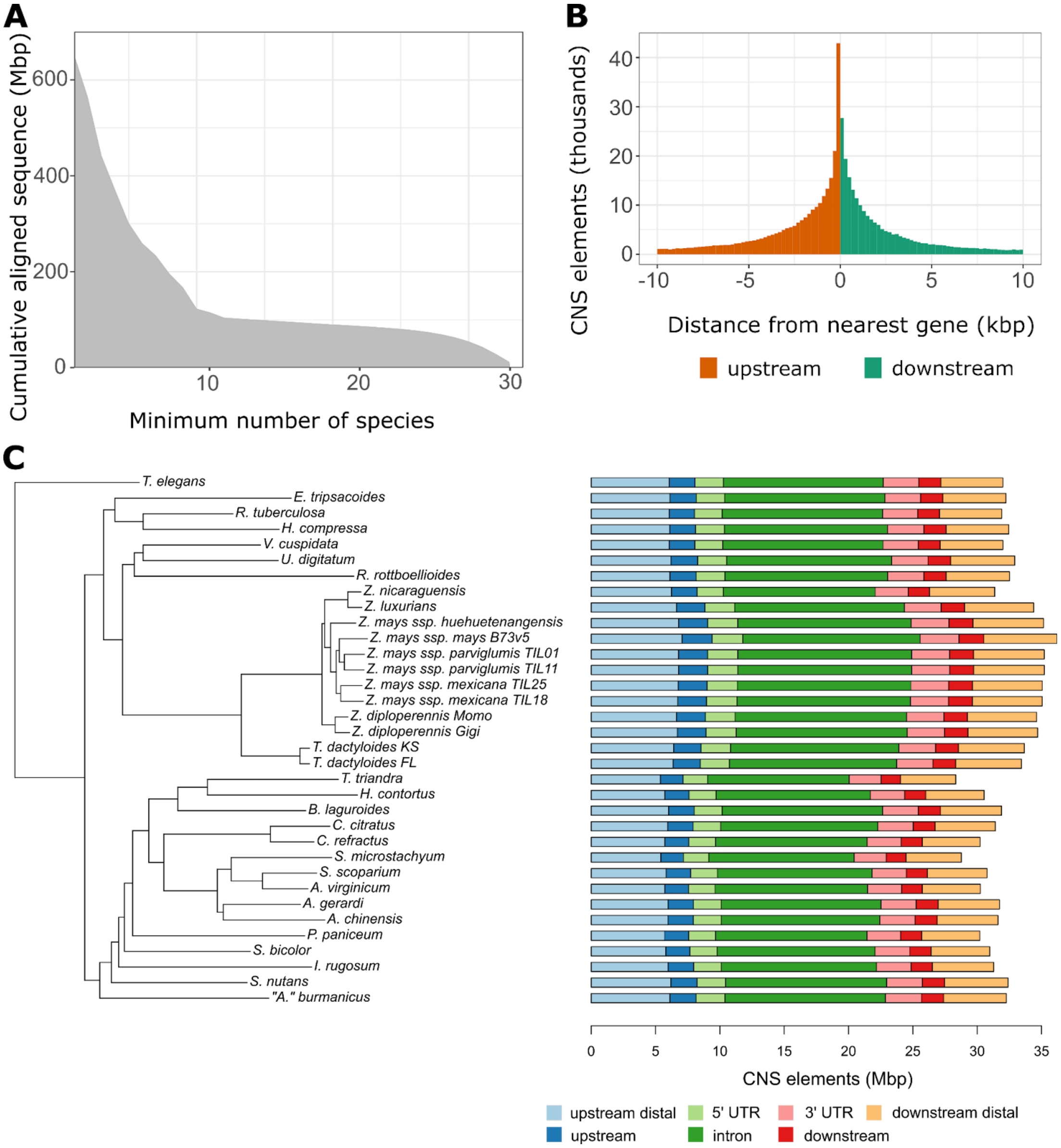

**Supplemental Figure S8. Gene-gene distance is high and bimodal in paleotetraploid Tripsacinae and *E. tripsacoides***

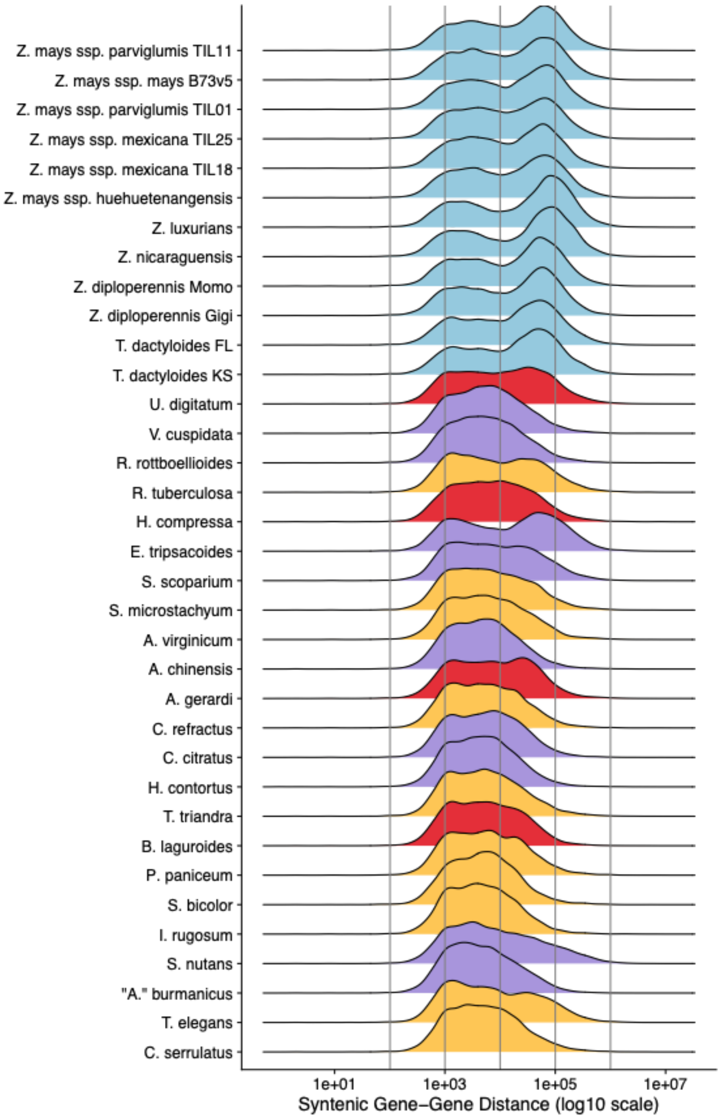
