## Supplemental Text for "Extensive genome evolution distinguishes maize within a stable tribe of grasses"

### *Heterozygosity, Life History, and Mating System*

Heterozygosity varies across our sample, and follows expectations based on mating system, life history, and range size, insofar as those are known. We measure the highest heterozygosity in *T. elegans* (5.1%), where flowers are chasmogamous (Thompson, 2019). Further, this monotypic genus is self-incompatible (Sisodia, 1970a), and populations segregate for translocations without a cost to pollen fertility, likely mediated by the formation of ring chromosomes during meiosis (Sisodia, 1970b, 1970a). The species is native to both Africa and the Indian subcontinent, and together a large range, obligate outcrossing, and cytological pairing likely contribute to this high observed heterozygosity. We also find high heterozygosity in *H. contortus* (4.8%), which is broadly distributed at latitudes from 35 degrees north to 35 degrees south (Emery & Brown, 1958). It is an aposporous apomict, a mode of reproduction that can lead to dominance of a single well-adapted genotype. Instead, studies of *H. contortus* have shown high diversity (Carino & Daehler, 1999; Chu et al., 2021). Further, seed dispersal is facilitated by a long awn that twists when wet or while drying, and dispersal can occur over long distances, with a majority of experimentally applied seeds still present on clothing after a five kilometer hike (Ansong & Pickering, 2013). We find high heterozygosity in another widely introduced species, *C. citratus* (lemongrass, 3.5%). This species is often vegetatively propagated, rarely producing seed (Oyen, 1999), so high heterozygosity may result from the establishment of successful clonal lineages that complement deleterious alleles.

Many Andropgoneae polyploids are apomictic, allowing favorable allelic combinations to accumulate in a heterozygous state, but limiting population-wide genetic diversity from clonal stands. Known apomicts we have sampled include *H. contortus*, described above, *T. triandra*, and *T. dactyloides*. Both *T. triandra* and *T. dactyloides* are mixed ploidy species, with diploid and polyploid individuals. In these species, apomixis has only been observed in polyploids, while diploids are sexual (Brown & Emery, 1957; Grimanelli et al., 1998). We observe heterozygosity of 1.7% in *T. triandra*. Our sampled *T. dactyloides* individuals are both diploids, and the difference in heterozygosity levels (1.2% for Kansas sample, 0.4% for Florida sample) appears due to differences in historical inbreeding.

Many Andropogoneae species are self-incompatible, due to the gametophytic grass S-Z incompatibility system (Baumann et al., 2000). Sometimes, this differs with ploidy level, which may impact observed heterozygosity. Polyploids of a congener to *H. compressa*, *Hemarthria altissima*, have been shown to undergo sexual reproduction (Wilms et al., 1970), and while diploids were self-incompatible, polyploids were self-compatible in bagged inflorescences. Our 2.1% heterozygosity for the *H. compressa* hexaploid is consistent with arising from outcrossing. Both *A. gerardi* and *S. nutans* (McKone et al., 1998) are self-incompatible, with abnormal pollen tube growth in self-pollinated pistils. *A. gerardi* shows relatively high heterozygosity (2.1%), but *S. nutans* shows lower heterozygosity (0.35%), while both have similar range size and life history.

Several species have been reported to be self-compatible, but our sampled individuals appear to arise from outcrossing instead of selfing. *B. laguroides* is self-compatible and not apomictic (Allred & Gould, 1983). Our sampled individual appears to arise from outcrossing, with 1.7% heterozygosity. However, we observed this individual producing self-pollinated seeds in the greenhouse, supporting that this individual is self-compatible. Australian species of *Schizachyrium* have been noted to predominately self due to cleistogamous flowers (Blake, 1974), but our sampled individuals of both *S. microstachyum* (0.40%) and *S. scoparium* (0.45%) have heterozygosity values more consistent with outcrossing.

Self-compatibility has been measured for several Andropogoneae taxa in which we observe very low heterozygosity, although reports are often summarized at the level of genus (Connor 1979). *Rottboellia exalta*, congener to *R. tuberculosa*, is self-compatible and commonly self-pollinated (Heslop-Harrison, 1959), although outcrossing may be elevated by longer development before exposure to short days. Our individual shows very low (0.00012%) heterozygosity, consistent with arising via self-fertilization. Other self-compatible taxa with very low heterozygosity (0.00022%) include *A. virginicum*, where flowers are cleistogamous, preventing outcrossing (Campbell, 1982). Our sample of annual teosintes (*Z. mays*) also show low heterozygosity, likely a result of the at least six generations of selfing to create inbred lines in the teosinte inbred lines (TILs) or small census population size of *Z. mays* subsp. *huehuetenangensis*. We are unaware of reports of self-compatibility in *U. digitatum*, which appears to arise via selfing (heterozygosity 0.00028%).

#### *Additional Cytological Evidence*

For many species, the chromosome count is consistent with an ancestral base chromosome number of n=10 (Supp. Table S3). We did not generate chromosome counts for all species, but records exist in the literature for some. These include *Rhytachne rottboellioides*, which has been noted to be a tetraploid with n=16 (Davidse and Pohl 1974), but see earlier estimates of 2n=36 (Gould and Soderstrom 1967). *Hemarthria compressa* from the same accession as our assembly (GRIN PI 404118) was identified to be 2n=54 (Quesenberry et al. 1982). *Chrysopogon serrulatus* occupies a wide range of ploidy, but our sample from Pakistan appears to be a diploid with 2n=20, as seen in this species in India (Saxena and Gupta 1969).

We noted that *Elionurus tripascoides* chromosomes are large and fluffy compared to other species (Supp Fig 1), with 2n=20. The large size of *Elionurus* chromosomes has been noted as a "striking feature" in comparative cytological surveys (Brown 1951), and the related species *Elyonurus argenteus* was noted to contain the largest known chromosomes in the Andropogoneae (Celarier 1957). Comparative cytology across two accessions of *Elionurus tripsacoides* suggested 2n=20, with different positions of the nucleolus organizer, and varying lengths of some chromosomes between accessions from Texas and Veracruz (Chandravadana and Galinat 1976). This could support fluidity in the chromosome complement. Earlier work supported n=5 for African *E. argenteus*, potentially with repeated fusions between terminal knobs complicating cytology (Celarier 1957). Mapping PacBio reads back to this assembly shows a heterozygosity of 0.0075, consistent with a haploid assembly of a tetraploid, and supporting 2n=4x=20.

Our *Rottboellia tuberculosa* assembly contains a single copy of all syntenic anchors, indicating a diploid. However, both flow cytometry (1,751 Mb) and total assembly size (1,572 Mb) are larger than other sampled diploids. We were unable to generate chromosome squashes for this individual, but closely related congener *Manisuris cylindrica* has been reported with 2n=18 (Brown 1951; Gould 1956). While we cannot exclude the possibility this individual is an autotetraploid with very low diversity, we categorize this individual as a diploid.

Although *Thelepogon elegans* has high diversity for a diploid, it has been shown cytologically to be diploid (2n=2x=10) with reduced chromosome number n=5 (Naithani & Sisodia, 1967; Sisodia, 1970b). However, some cells of this individual contained 2n=20 chromosomes, and authors suggested this syndiploidy results from failed reduction after mitosis and may be a mechanism for polyploid formation (Sisodia, 1970b). The potential for ring chromosomes and possible permanent translocation heterozygosity (Sisodia, 1970b) may be responsible for the elevated diversity we observe.
